# Probing function in ligand-gated ion channels without measuring ion transport

**DOI:** 10.1101/2022.03.08.483473

**Authors:** Nicole E. Godellas, Claudio Grosman

**Affiliations:** Department of Molecular and Integrative Physiology, University of Illinois at Urbana-Champaign; Center for Biophysics and Quantitative Biology, University of Illinois at Urbana-Champaign; Neuroscience Program, University of Illinois at Urbana-Champaign

**Author notes:** To whom correspondence should be addressed. E-mail address; ORCID identifier: 0000-0001-8093-8362; Postal address: 407 S. Goodwin Ave., 524 Burrill Hall, Urbana, IL 61801, USA.

## Abstract

Although the functional properties of ion channels are most accurately assessed using electrophysiological approaches, a number of experimental situations call for alternative methods. Here, working on members of the pentameric ligand-gated ion-channel (pLGIC) superfamily, we focused on the practical implementation of, and the interpretation of results from, equilibrium-type ligand-binding assays. Ligand-binding studies of pLGICs are—by no means—new, but the lack of uniformity in published protocols, large disparities between the results obtained for a given parameter by different groups, and a general disregard for constraints placed on the experimental observations by simple theoretical considerations suggested that a thorough analysis of this classic technique was in order. To this end, we present a detailed practical and theoretical study of this type of assay using radiolabeled α-bungarotoxin, unlabeled small-molecule cholinergic ligands, the human homomeric α7-AChR, and extensive calculations in the framework of a realistic five-binding-site reaction scheme. Furthermore, we show examples of the practical application of this method to tackle two long-standing questions in the field: whether ligand-binding affinities are sensitive to binding-site occupancy, and whether mutations to amino-acid residues in the transmembrane domain can affect the channel’s affinities for ligands that bind to the extracellular domain.

## INTRODUCTION

Regardless of structural differences between superfamilies, all neurotransmitter-gated ion channels (NGICs) are integral membrane proteins formed by, essentially, two modules: an extracellular domain (ECD) that harbors the neurotransmitter-binding (“orthosteric”) sites, and a transmembrane domain (TMD) that forms the transmembrane aqueous pore. Conformational changes in the ECD result in different affinities for the neurotransmitter (“low” and “high”), whereas conformational changes in the TMD result in pores that either conduct (“open”) or do not conduct (“closed” and “desensitized”) ions. These conformational changes are not independent of each other, but rather, are thought to be strictly correlated (“coupled”) in such a way that conformations of the receptor-channel that bind neurotransmitter with low affinity have non-conductive closed pores, whereas those that bind neurotransmitter with high affinity have either ion-conductive open pores or non-conductive desensitized pores (Chang and Weiss, 1999; Grosman and Auerbach, 2001; Jackson, 1989). Thus, the interconversion of these ligand-gated ion channels between the closed, open, and desensitized states (hereafter, “gating”) can be inferred by measuring the transport of ions through their pores or by estimating the extent of ligand binding, that is, by following the operation of one module or the other.

The higher sensitivity and time resolution of methods that measure ion transport— particularly, those that measure the associated ion currents—easily explain their dominance over ligand-binding studies as experimental approaches to probe function in NGICs. However, one can imagine a variety of circumstances under which the measurement of currents or ion fluxes is not possible or not desired: a) Mutations may render an ion channel electrically silent by stabilizing non-conductive conformations or greatly reducing the single-channel conductance; b) An agonist may desensitize a channel too quickly; c) Studies of the interaction between ion permeation and gating (“permeation–gating coupling”) may require that function also be studied in the absence of ion flow; d) Maneuvers that modify the lipid composition of the plasma membrane may render the formation of high-resistance patch-clamp seals unlikely and vesicles for ion-flux assays, leaky; e) Studies of the effect of the lipid environment on function may require that a channel be solubilized in detergent micelles so as to establish a baseline behavior; and f) Comparative studies of the effect of different types of membrane mimetic in the context of structural-biology efforts may require that channel function be studied in lipid nanodiscs.

Furthermore, even if current measurements were possible, low single-channel conductance, poor expression levels, and/or hard-to-control time-dependent changes in channel activity upon patch-clamp seal formation or excision (usually referred to as “run-down” or “run-up”) may render electrophysiological studies highly impractical; in these cases, ligand-binding assays provide a robust alternative.

Ligand-binding experiments often take the form of concentration–response assays in which some direct or indirect measure of binding is recorded and plotted against the concentration of ligand. The resulting curves are fitted with empirical functions—most commonly, a “Hill equation”—and the values of the estimated parameters (that is, a half-effective concentration and a “Hill coefficient”) are used to characterize the receptor–ligand complex under different experimental conditions. Although, with some exceptions, these empirical parameters cannot be expressed easily in terms of the underlying equilibrium constants of state interconversions, their use is favored because they are convenient. Fitting the observations with more realistic, mechanism-based equations (having many more parameters) would likely be impossible (e.g. Hines et al., 2014).

In the context of NGICs, ligand-binding studies have been mostly applied to the members of the pentameric ligand-gated ion-channel (pLGIC) superfamily (also known as “Cys-loop receptors), and within these, to the muscle-type nicotinic acetylcholine receptor (AChR) (Blount and Merlie, 1989, 1988; Covarrubias et al., 1986; Franklin and Potter, 1972; Fulpius et al., 1972; Maelicke et al., 1977; Quast et al., 1978; Sine and Taylor, 1979; Weber and Changeux, 1974a, 1974b, 1974c; Weiland and Taylor, 1979) and the α7 AChR (Corringer et al., 1995; Gopalakrishnan et al., 1995; Peng et al., 1994). Undoubtedly, this is because of the availability of a powerful tool: α-bungarotoxin (α-BgTx; (Lee, 1970)), a 74-amino-acid snake toxin that binds to muscle-type and α7 AChRs competitively with orthosteric ligands (that is, ACh, nicotine, and their analogs) and dissociates from the complex slow enough to allow the effective physical separation of bound from unbound label. However, although these assays have been in use since the 1970’s, we noticed a lack of uniformity in published protocols, as well as large discrepancies in the values of parameters estimated from even the simplest type of experiments. For example, for the chicken α7-AChR [in the context of an ECD–TMD chimera having the rat serotonin-receptor type 3A’s (5-HT_3A_R’s) TMD], values of the α-BgTx dissociation equilibrium constant (*K*_D_) from the closed-channel conformation of 70 pM (Rangwala et al., 1997) and 4.2 nM (Pittel et al., 2010) have been reported for receptors on resuspended cells. Similarly, for the wild-type human α7-AChR, values of this *K*_D_ have been reported to be 0.8 nM (Peng et al., 1994) and 4 nM (Tillman et al., 2016) in detergent-solubilized receptors, 0.7 nM (Gopalakrishnan et al., 1995) and 7 nM (Peng et al., 2005) in resuspended membrane homogenates, and 1.2 nM (Shabbir et al., 2021), 2.7 nM (daCosta et al., 2015) and 26 nM (Sine et al., 2019) in resuspended cells. Moreover, values of the Hill coefficient of small-molecule ligands reported in the literature of these ion channels often run counter theoretical expectations, with values significantly different from unity for antagonists, and values significantly lower than unity for agonists.

Here, we set out to optimize the values of the different experimental variables in otherwise “classic” concentration–response assays at equilibrium. To this end, we used radio-iodinated α-BgTx, small-molecule cholinergic ligands, and the full-length homomeric α7-AChR expressed in HEK-293 cells. Furthermore, using a realistic five-binding-site reaction scheme, we investigated the quantitative relationships between the empirical parameters estimated from equilibrium concentration–response curves and the underlying equilibrium constants of ligand-binding and ion-channel gating. Finally, we end the paper with examples of the practical application of this method to tackle two long-standing questions in the field: whether the ligand-binding affinity for any of the orthosteric sites is sensitive to the occupancy of the other orthosteric sites, and whether mutations to amino-acid residues in the TMD can affect the channel’s affinities for ligands that bind to the ECD.

## MATERIALS AND METHODS

### cDNA clones, mutagenesis, and heterologous expression

Complementary DNA (cDNA) coding the human α7-AChR (UniProt accession number: P36544) in pcDNA3.1 was purchased from addgene (#62276); cDNA coding isoform 1 of human RIC-3 (accession number: Q7Z5B4; (Treinin, 2008)) in pcDNA3.1 was provided by W. N. Green (University of Chicago, IL); cDNA coding human NACHO (TMEM35A; accession number: Q53FP2; (Gu et al., 2016)) in pCMV6-XL5 was purchased from OriGene Technologies Inc. (#SC112910); cDNA coding the “cut-and-splice” (“CS”) chimera between the ECD of the α7-AChR from chicken and the TMD of β-GluCl from *C. elegans* in pMT3 (Cymes and Grosman, 2021) was obtained by mutagenesis of a related clone provided by Y. Paas (Bar-Ilan University, Israel) (Sunesen et al., 2006); cDNA coding the human–*C. elegans* counterpart of the chicken–*C. elegans* CS α7-AChR–β-GluCl chimera was obtained by mutagenesis (the mutations were: L34V, T56S, M60L, Y81S, N99T, L106Q, K163H, N165K, T172S, S192P, T206S, S208R, and I220V); cDNA coding the human acid-sensing ion channel subunit 1 (ASIC1; accession number: P78348) in pCR-BluntII-TOPO was purchased from horizon (#MHS6278-211689646) and was subcloned in pcDNA3.1; and cDNAs coding the mouse β1, δ, and ε subunits of the (muscle) AChR (accession numbers: P09690, P02716, and P20782, respectively) in pRBG4 were provided by S. M. Sine (Mayo Clinic, Rochester, MN). Mutations were engineered using the QuikChange kit (Agilent Technologies), and the sequences of the resulting cDNAs were verified by dideoxy sequencing of the entire coding region (ACGT). Wild-type and mutant channels were heterologously expressed in transiently transfected adherent HEK-293 cells grown at 37°C and 5% CO_2_. cDNAs coding the human α7-AChR, human RIC-3, and human NACHO were co-transfected using 125, 687.5, and 687.5 ng cDNA/cm^2^, respectively; cDNAs coding the chicken–*C. elegans* α7-AChR–β-GluCl chimera, the human–*C. elegans* α7-AChR–β-GluCl chimera or human ASIC1 were transfected using 187.5 ng cDNA/cm^2^; and cDNAs coding the mouse β1-, δ-, and ε-AChR subunits were co-transfected using 62.5 ng cDNA/cm^2^ each. Transfections were performed using a calcium-phosphate-precipitation method and proceeded for 16–18 hr after which the cell-culture medium (DMEM; Gibco) containing the DNA precipitate was replaced by fresh medium. As a control of the non-specific binding of α-BgTx to cells, HEK-293 cells were transiently transfected with cDNA coding the human ASIC1 or the mouse β1-, δ-, and ε-AChR subunits.

These cells were incubated with [^125^I]-α-BgTx (in the absence of unlabeled competitive ligand) under the same conditions as were the cells expressing wild-type or mutant α7-AChRs. The resulting non-specific-binding values were used to calculate specific binding for the saturation curves, and for the subset of competition curves in which the highest concentrations of unlabeled ligand were unable to displace the specifically bound [^125^I]-α-BgTx completely.

### Ligand-binding assays

24-h after changing the cell-culture medium, transfected cells were resuspended in a Hepes-buffered sodium-saline solution (in mM, 142 NaCl, 5.4 KCl, 1.8 CaCl_2_, 1.7 MgCl_2_, and 10 Hepes/NaOH, pH 7.4) by gentle agitation, and divided in 1-mL aliquots in 1.7-mL plastic tubes. Ligand binding-reaction mixtures were incubated at the indicated temperature and for the indicated duration with constant rotation, and upon completion, cell-bound label was separated from unbound label by centrifugation at 16,000 × *g* for 3 min at room temperature. In order to reduce the amount of non-specifically-bound label, these pellets were resuspended in 1-mL of Dulbecco’s phosphate-buffered saline (pH 7.4; Gibco), vortexed for 30 s, and pelleted again at 16,000 × *g* for 3 min at room temperature; this resuspension–pelleting procedure was repeated twice. Finally, the washed pellets were resuspended in a solution containing 0.1 N NaOH and 1% (w/v) sodium dodecyl sulfate (SDS), and incubated at 65–70°C for 30 min. The radioactivity and protein content of each solubilized pellet were estimated: ^125^I radioactivity was measured using a Wiper^TM^ 100 γ-counter (Laboratory Technologies, Inc.) that we calibrated (efficiency = 0.826) using a QCI-501 standard (Reflex Industries), and the amount of protein was measured using the bicinchoninic-acid assay (BCA; ThermoFisher) and a freshly prepared bovine serum-albumin (ThermoFisher) calibration curve. The number of transfected cells contained in each reaction tube of any given curve was adjusted, by trial and error, so as to minimize the depletion of labeled and unlabeled ligands while ensuring high-enough a signal. For some constructs, the expression of receptors was so high that the amount of transfected cells that satisfied this criterion resulted in pellets that were too small to handle reliably. In these cases, to increase the size of the pellets, transfected cells were mixed with non-transfected cells. Most experiments were repeated several times, each one using two replicates per concentration of [^125^I]-α-BgTx (in saturation experiments) or unlabeled ligand (in competition experiments). All curves corresponding to a given set of conditions were fitted globally with a Hill equation using SigmaPlot 14 (Systat Software Products). For display purposes, these data points were normalized using the globally fitted parameters, averaged, and plotted as mean ± 1 standard error (SE) of the several replicates. For all fits, the reciprocal of the *y*-axis variable was used as weight, and parameter SEs were computed using the reduced χ^2^ statistic. [^125^I]-α-BgTx was purchased from PerkinElmer (initial specific activity ≅ 80–140 Ci/mmol); MLA and DHβE, from Tocris Bioscience; and carbamylcholine, choline, and nicotine, from MilliporeSigma.

## RESULTS

### An overview of ligand-binding assays

Ligand-binding assays often entail the use of labeled ligands that allow the direct estimation of the number of ligand-molecules bound. When the characterization of the interaction between a labeled ligand and its receptor is of interest, the experiment takes its simplest form: receptor and ligand are incubated in mixtures containing an approximately constant concentration of receptors and a variable concentration of ligand ranging from zero (or very small, if the curves are displayed on a logarithmic *x* axis) to saturating. The binding reactions are allowed to proceed until equilibrium is attained, and then, the label associated to ligand–receptor complexes is measured. Here, we will refer to these assays as “saturation-binding” assays. However, it may also be of interest to characterize the interaction between the receptor in question and other ligands that may not be readily available in labeled form, but that may bind to the receptor in a manner that is mutually exclusive with the binding of the available labeled ligand. In this case, two alternative (seemingly similar, but conceptually very different) approaches can be taken (Weber and Changeux, 1974a). In one of them, mixtures containing a fixed concentration of receptor and a range of concentrations of unlabeled ligand are incubated until equilibrium between the two is reached. Then, in a second step, a fixed concentration of the labeled ligand is added to each reaction, and the amount of binding is recorded as a function of time. The extent to which the initial rate of labeled-ligand binding is slowed down with increasing concentrations of unlabeled ligand is then plotted and analyzed. In this method, the purpose of the labeled ligand is to act as a mere reporter of the number of sites left unoccupied by the equilibrated mixture of unlabeled ligand and receptor. The binding of the labeled ligand is analyzed over very short times, much shorter than needed for equilibrium to be attained by the three components of the mixture. These “kinetic” studies are typically referred to as “protection” assays. The alternative approach consists of the incubation of receptor (at a fixed concentration), labeled ligand (also, at a fixed concentration), and unlabeled ligand (at a variable concentration) until equilibrium between all three components of the ternary mixture is reached; only then is the amount of receptor-bound label measured. We will refer to this method as the “equilibrium binding-competition” approach. The latter is the method we favor as a probe for pLGIC function when ion transport is not measured, and the method we elaborate on, below.

### The equilibrium binding-competition approach

In the presence of two ligands that bind in a mutually exclusive manner (such as the labeled and unlabeled ligands), a homopentameric pLGIC with five identical binding sites can exist in several different ligation and conformational states. These states are indicated in the reaction scheme in Figure 1, which (in keeping with experimental observations (Grosman and Auerbach, 2001, 2000a, 2000b; Jackson, 1984)), is built around the concepts of unliganded gating and binding–gating thermodynamic cycles (Jackson, 1989). For the sake of clarity of display and simplicity of calculation, the open and desensitized conformations—that is, the conformations of the channel that bind agonists with higher affinity—were lumped together and are denoted, collectively, as “O” or “open”.

**Figure 1.**
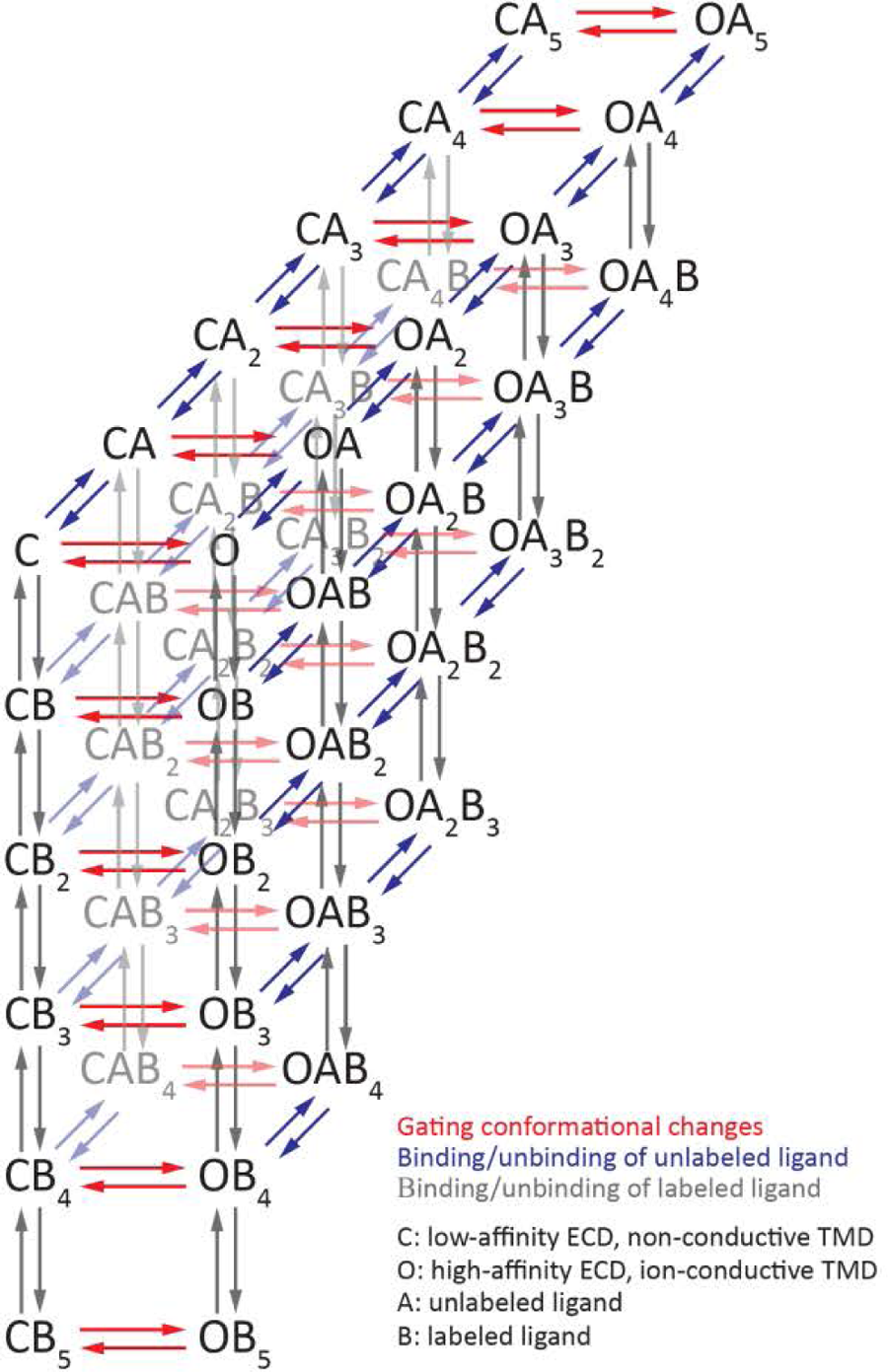
A theoretical framework. A 42-state reaction scheme used to calculate equilibrium ligand-binding concentration–response curves of homopentameric pLGICs as they interact with binary mixtures of ligands competing for the five orthosteric neurotransmitter-binding sites. Calculated curves were used to learn about the behavior of the system and guide the interpretation of the experimentally obtained curves. The reaction scheme is composed of elementary thermodynamic cycles consisting of gating conformational changes and ligand-binding steps; thus, the values of the equilibrium constants within each cycle were constrained by detailed balance. The scheme assumes that all five binding sites are identical. Unless otherwise stated, our calculations also assumed that the affinities for labeled and unlabeled ligands are independent of receptor occupancy; affinities only depend on whether the receptor-channel is closed or open. For the sake of simplicity, the open and desensitized conformations— that is, the conformations of the channel that bind neurotransmitter and other agonists with higher affinity—were lumped together, not only for the graphical display of this reaction scheme, but also, for the calculations. Thus, in the context of this paper, “open state” refers to both, open and desensitized states. Throughout this study, the following symbols were used. *K_C_*_⇌_*_O_*: gating equilibrium constant of the unliganded channel; *K_DA,closed_* and *K_DA,open_*: dissociation equilibrium constants of unlabeled ligand from the closed and open states, respectively; and *K_DB,closed_* and *K_DB,open_*: dissociation equilibrium constants of labeled ligand from the closed and open states, respectively.

One of the key advantages of methods that use labeled ligands is the most straightforward relationship between the observed signal and the ligand-binding phenomenon under study. Indeed, upon subtraction of non-specific binding, the signal is directly proportional to the mean number of binding sites per receptor occupied by the labeled ligand. In turn, the expected value of the latter quantity as a function of the concentration of ligand (whether labeled or unlabeled) in concentration–response curves can be calculated for any set of equilibrium constants. Figure 2 shows some examples of these calculations for a variety of parameters in a hypothetical competition assay in the context of the reaction scheme in Figure 1.

**Figure 2.**
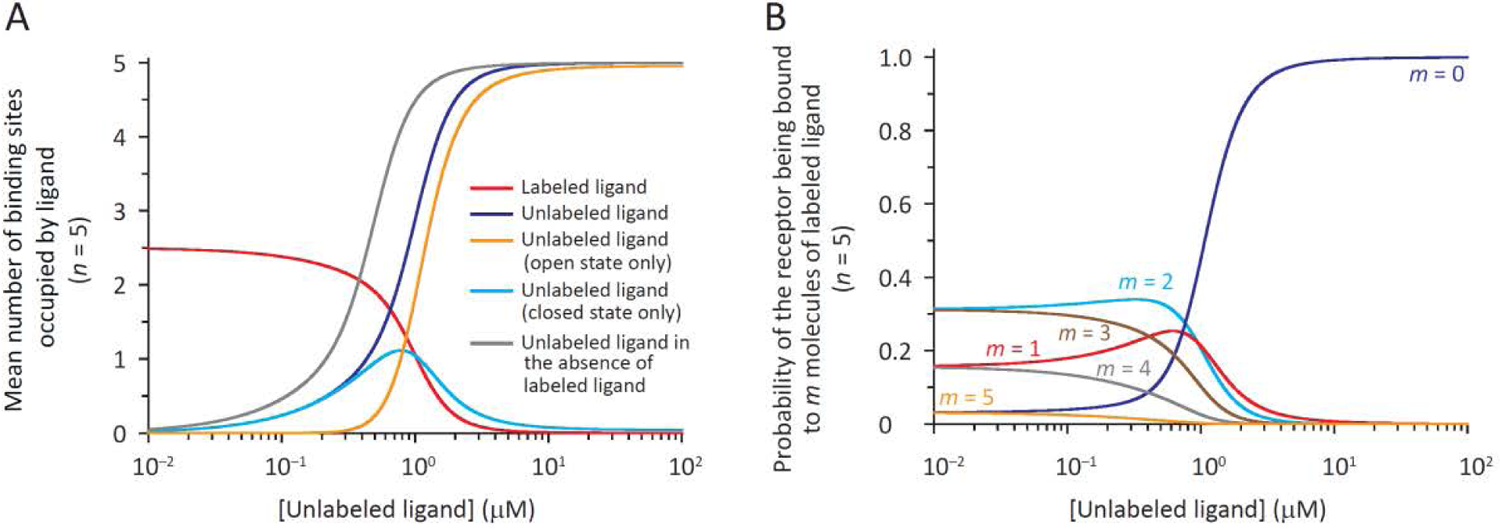
Calculated concentration–response curves and various probabilities. (**A**, **B**) Various quantities were calculated on the basis of the reaction scheme in Figure 1 for a hypothetical competition experiment between a labeled ligand and an unlabeled ligand. The parameters were: *K_C_*_⇌_*_O_* = 10^-7^; *K_DA,closed_* = 1 μM; *K_DA,open_* = 15 nM; *K_DB,closed_* = 1 nM; and *K_DB,open_* = 4 nM. Thus, the unlabeled ligand was assumed to be an agonist (like ACh, nicotine or carbamylcholine) and the labeled ligand, a weak inverse agonist (like α-BgTx; (Bertrand et al., 1997; Jackson, 1984). The total number of binding sites (*n*) was 5. The principle of detailed balance and the notion that the binding sites are identical and independent were applied to calculate the gating equilibrium constants of the channel in its different ligation states. The ratio between the fixed and half-saturation concentrations of labeled ligand was unity, and hence, the mean number of binding sites per receptor occupied by labeled ligand at zero concentration of unlabeled ligand was one-half of their total number. Only the binding of labeled ligand (red plot in *A*) can be estimated experimentally. In *A*, the plot in blue is the sum of those in orange and cyan. Also in *A*, the plot in grey shows what a binding curve of the unlabeled ligand would look like if a competing ligand were not used in the assay. For both panels, the concentration of unlabeled ligand (on the *x*-axes) corresponds to the concentration of unbound (“free”) unlabeled ligand at equilibrium.

### Effects of the labeled ligand on the observations

In equilibrium binding-competition assays, the experimenter is usually interested in elucidating properties of the receptor as it interacts with the unlabeled ligand—the labeled ligand acts as a probe. However, because the binding of both types of ligand is expected to reach equilibrium before the signal is recorded, the (fixed) concentration of the labeled ligand and its affinities for the different conformational states of the receptor-channel are expected to affect the steepness and displacement along the *x*-axis of competition concentration–response curves. In other words, rather than being a mere reporter, the labeled ligand is an integral part of the binding reactions with potential effects on the results.

Labeled ligands used in competition assays are often antagonists (that is, molecules that bind with indistinguishable affinities to the different conformations of the receptor-channel) or inverse agonists (molecules that bind with higher affinity to the “resting”, closed conformation), such as α-BgTx, in the case of AChRs, and strychnine, in the case of glycine receptors. Moreover, most wild-type pLGICs almost exclusively populate the low-affinity closed conformation when unliganded. Quite conveniently, it can be shown that, under these conditions, competition curves remain essentially unaffected by the properties of the labeled ligand as long as the ratio between its fixed concentration used in the assay and the concentration that half-saturates the receptor remains constant. Thus, because mutations can affect the affinities of the receptor for not only unlabeled ligands, but also, for the labeled ligand, it becomes necessary to characterize the interaction between each new construct and the latter before competition assays can be performed. Although some authors have used labeled agonists in competition assays (such as radiolabeled epibatidine in studies of the ACh-binding protein; (Kaczanowska et al., 2014)), in this paper, we restricted our analysis to labeled ligands that act as antagonists or inverse agonists.

Figure 3 illustrates these concepts with concentration–response curves calculated in the context of the reaction scheme in Figure 1 for several hypothetical situations (denoted *i*–*viii*). For all of them, the closed ⇌ open, gating equilibrium constant of the unliganded channel was assumed to be 10^-7^, in keeping with experimental estimates of this quantity for the wild-type muscle AChR (Jackson, 1984; Purohit and Auerbach, 2009) and the known low unliganded activity of most other wild-type pLGICs. In *i* (Fig. 3, A and B), the dissociation equilibrium constant of the labeled ligand from the closed state was 1 nM and that from the open state was 4 nM (and thus, from detailed balance, the gating equilibrium constant of the channel bound to five molecules of this weak inverse agonist was 10^-7^ × (1/4)^5^ ≅ 10^-10^). Furthermore, the dissociation equilibrium constant of the unlabeled ligand from the closed state was 1 μM, and that from the open state was 15 nM (and thus, the gating equilibrium constant of the channel fully bound to this strong agonist was 10^-7^ × (1/0.015)^5^ ≅ 132). With these values, it can be calculated that the concentration of labeled ligand that half-saturates the receptor is ∼1 nM, and the fixed concentration of labeled ligand used to calculate the competition curve was chosen to also be 1 nM—in this way, the ratio between these quantities was unity. Under these conditions, the concentration of unlabeled ligand that displaces half of the bound labeled ligand can be calculated to be ∼0.88 μM. In *ii*, we modeled a perturbation (say, a mutation) that increases the dissociation equilibrium constants of the unlabeled ligand from the closed and open states by a factor of 100 (the same factor for both) without affecting the dissociation equilibrium constants of the labeled ligand. Therefore, the concentration of labeled ligand that half-saturates the receptor remains ∼1 nM, and the fixed concentration of labeled ligand used in the calculation was again chosen to be 1 nM. Under these conditions, the concentration of unlabeled ligand that displaces half of the bound labeled ligand can be calculated to also increase by a factor of ∼100: it is ∼88 μM. In *iii*, the hypothetical perturbation decreases the dissociation equilibrium constants of the labeled ligand from the closed and open states by a factor of 10 in addition to increasing the dissociation equilibrium constants of the unlabeled ligand by a factor of 100, as in *ii*. With these values, it can be calculated that the concentration of labeled ligand that half-saturates the receptor is ∼100 pM. Assuming that the experimenter performed a saturation curve and noted this change, the fixed concentration of labeled ligand used in the calculation of the competition curve was adjusted to 100 pM, so as to keep a constant ratio between the fixed and half-saturation concentrations across constructs. Under these conditions, the concentration of unlabeled ligand that displaces half of the bound labeled ligand can be calculated to also be ∼88 μM. That is, provided that changes in the affinity of the receptor for the labeled ligand are detected and accounted for, they have no effect on the binding-competition curves. Finally, in *iv*, the situation in *iii* is illustrated assuming that the experimenter did not notice the change in affinity for the labeled ligand, and thus, still used a concentration of 1 nM of it throughout the assay. Under these conditions, the concentration of unlabeled ligand that displaces half of the bound labeled ligand can be calculated to be quite larger: ∼490 μM. Indeed, at a concentration of 1 nM, a ligand with a 100-pM dissociation equilibrium constant would bind to ∼91% of the binding sites (∼4.5 out of 5 sites) rather than to only 50% of them. Figure 3 B shows the plots in Figure 3 A normalized to their respective maximum values.

**Figure 3.**
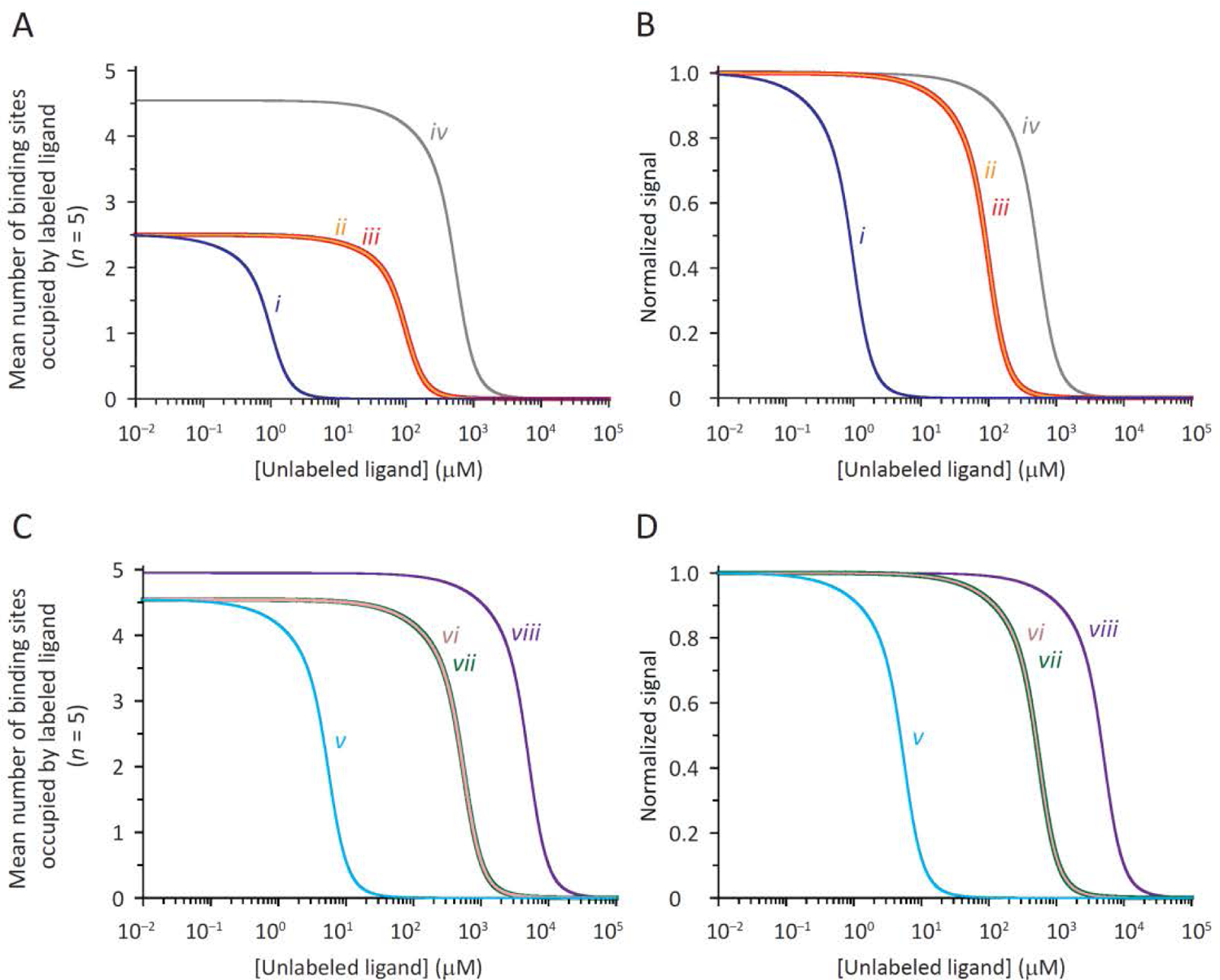
The importance of characterizing the interaction between the receptor and the labeled ligand. Calculated absolute and normalized competition concentration–response curves for four hypothetical scenarios involving perturbations that affect the affinities of the receptor for the unlabeled and labeled ligands. The parameters used to calculate the curves are indicated in detail in the text. (**A**, **B**) The plot in *i* represents a “baseline” curve. The ratio between the fixed and half-saturation concentrations of labeled ligand was unity. In *ii*, both the *K_DA,closed_* and *K_DA,open_* values of the hypothetical receptor-channel were assumed to increase (by a factor of 100), and thus, the corresponding curve shifted to the right. The ratio between the fixed and half-saturation concentrations of labeled ligand was unity. In *iii*, not only did the affinities for the unlabeled ligand change (exactly as in *ii*), but also did those for the labeled ligand: both *K_DB,closed_* and *K_DB,open_* decreased (by a factor of 10). However, because the latter change was assumed to be known, the fixed concentration of labeled ligand was adjusted so as to maintain a value of unity for the ratio between the fixed and half-saturation concentrations. As a result, curves *ii* and *iii* overlap completely. In *iv*, the change in labeled-ligand affinities was assumed to have gone unnoticed, and thus, the labeled-ligand fixed/half-saturation concentration ratio was no longer unity. (**C**, **D**) The plots in *v*–*viii* are the counterparts of those in *i*–*iv* (*A* and *B*) with a ratio of fixed-to-half-saturation concentrations of labeled ligand equal to 10 for the baseline condition. For all four panels, the concentration of unlabeled ligand (on the *x*-axes) corresponds to the concentration of unbound unlabeled ligand at equilibrium. In *iii* and *vii*, the curves are plotted with thicker lines to clearly show the *ii–iii* and *vi–vii* complete overlap.

Figure 3, C and D show the four situations depicted in Figure 3, A and B using a larger, by a factor of 10, concentration of labeled ligand. Although each curve in Figure 3, C and D is shifted to the right relative to its counterpart in Figure 3, A and B, keeping a constant ratio between the fixed and half-saturation concentrations of labeled ligand ensured that plots *vi* and *vii* are identical, much like plots *ii* and *iii* are.

As shown above with a few examples, the ratio between the fixed concentration of labeled ligand used in competition assays and the concentration of labeled ligand that half-saturates the receptor need not be unity. However, this ratio needs to be kept constant across constructs for comparisons between receptors that display different affinities for the labeled ligand to only reflect changes in the properties of the unlabeled ligand. Furthermore, for any given construct, this ratio needs to remain constant for comparisons across different experimental conditions, different competing unlabeled ligands, and different laboratories to be meaningful.

### Fitting the observations with empirical functions

Saturation and competition curves are often fitted with Hill equations, and the estimated parameter values are compared across experimental conditions. When normalized to the maximum signal bound, one-component Hill equations are fully characterized by two free parameters: a concentration (a half-saturation or a half-competition value, depending on the type of assay) and a Hill coefficient (*n_H_*). For saturation experiments:

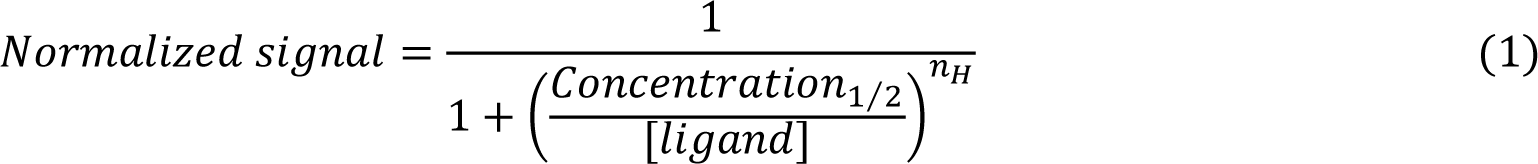

and, for competition experiments:

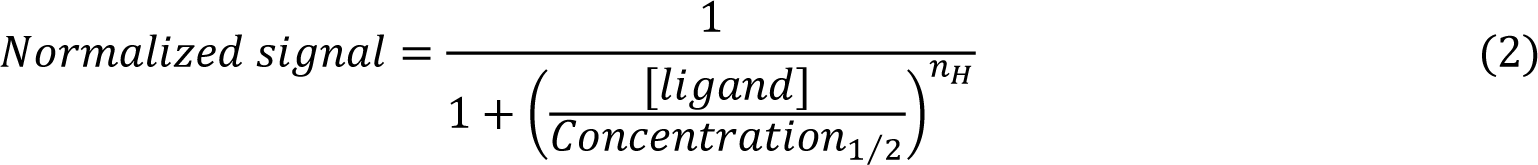

The numerical values of these two parameters depend on the equilibrium constants of all the underlying state-interconversion steps (both ligand-binding and gating), and in most cases, the mathematical relationship between them is far from straightforward. Furthermore, different combinations of values of the equilibrium constants of the different steps may result in similar values of the fitted parameters. As a result, assigning observed changes in binding-curve parameters to changes in the equilibrium constants of specific reaction steps is usually not possible. An exception to this generalization occurs when both the labeled and unlabeled ligands are antagonists. Certainly, in this case, it can be shown that the mean number of binding sites occupied by labeled ligand (*N*) is given by a simple expression that does not depend on the values of the gating equilibrium constants (which, for antagonists, are the same whether the channel is unliganded or ligand-bound):

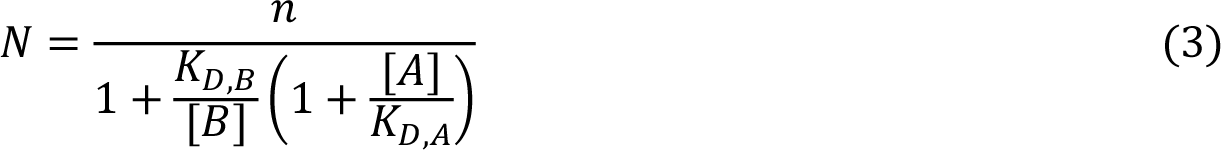

where, *n* denotes the total number of binding sites per receptor, and in keeping with the symbols in Figure 1, [*A*] and [*B*] denote the concentrations of unbound unlabeled and labeled ligands at equilibrium, respectively, and *K_D,A_* and *K_D,B_* denote the dissociation equilibrium constants of unlabeled and labeled ligands, respectively. In the particular case that the *K_D,B_*/[B] ratio is chosen to be 1, as was the case for our experiments, *Equation 3* can be rearranged to:

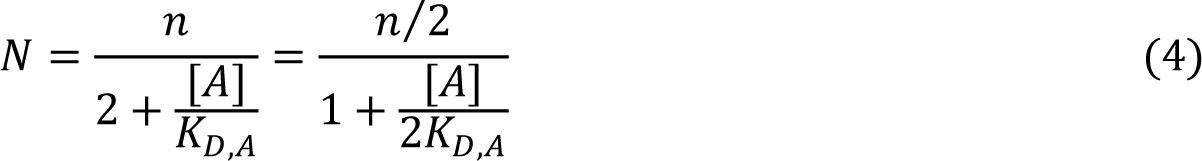

The value of *K_D,B_* can be estimated directly from the fitting of saturation curves (that is, from assays in which [*A*] = 0 for all points of the curve, and [*B*] is the variable). The corresponding expression of *N* follows from *Equation 3*:

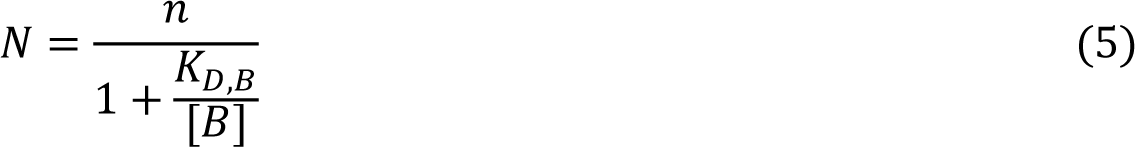

Thus, in competition experiments in which the *K_D,B_*/[*B*] ratio is equal to 1 (*Equation 4*), *N* decreases from *n*/2 (at [*A*] = 0) to zero, as [*A*] increases, and the value of [*A*] that displaces one-half of the bound labeled ligand (and thus, leaves one-fourth of the binding sites bound to the label) is numerically equal to 2×*K_D,A_*. More generally, the half-competition concentration is equal to *K_D,A_*×(1 + [*B*]/*K_D,B_*). It should be emphasized that these simple mathematical relationships are only accurate for antagonists competing against antagonists. In many cases, however, the unlabeled ligand is an agonist, and the counterparts of Eqs. 2–5 become more complicated because the various gating equilibrium constants no longer cancel. In these cases, neither ligand-dissociation nor gating equilibrium constants can be estimated directly from fits to ligand-binding curves.

As can be appreciated from a comparison of empirical *Equations 1 and 2* with mechanism-based *Equations 4 and 5*, Hill equations (with *n_H_* = 1) provide an accurate description of concentration–response curves only when the ligands involved are antagonists, regardless of the number of binding sites on the receptor (or, for all types of ligand, in the trivial case of receptors with a single binding site). In all other cases, Hill equations are only convenient approximations.

### Mechanistic interpretation of empirical parameters

In ligand-gated ion channels at equilibrium, binding and gating can be thought of as the elementary steps of thermodynamic cycles. Indeed, in the reaction scheme in Figure 1, each “square” containing “C” and “O” states is one such cycle. Therefore, because the product of equilibrium constants around a cycle must be unity, liganded gating can be considered to be determined by the unliganded-gating equilibrium constant, and the affinities of ligands for the closed and open states. In other words, the latter three quantities can be regarded as the independent variables (in the mathematical sense, at least) that, together, determine the values of the liganded-gating equilibrium constants. Assuming that the binding sites are identical and independent of each other:

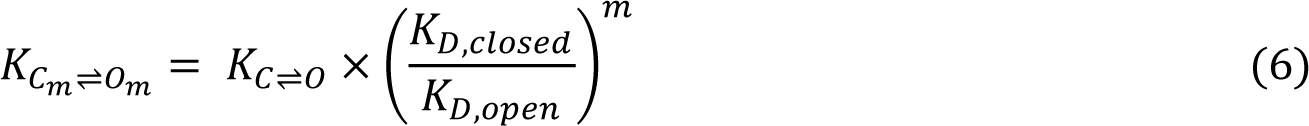

where *K_Cm_*_⇌_*_Om_* and *K_C_*_⇌_*_O_* denote the closed ⇌ open gating equilibrium constants of the liganded and unliganded receptors, respectively, *m* denotes the number of ligand molecules bound per receptor (1 ≤ *m* ≤ *n*), and *K_D,closed_* and *K_D,open_* denote the dissociation equilibrium constants of the ligand from the closed and open states, respectively. To learn about the relationship between the values of the empirical parameters in competition concentration–response curves and the values of the equilibrium constants of the “independent” variables, we calculated four hypothetical scenarios in the context of the reaction scheme in Figure 1 (Figs. 4–6).

**Figure 4.**
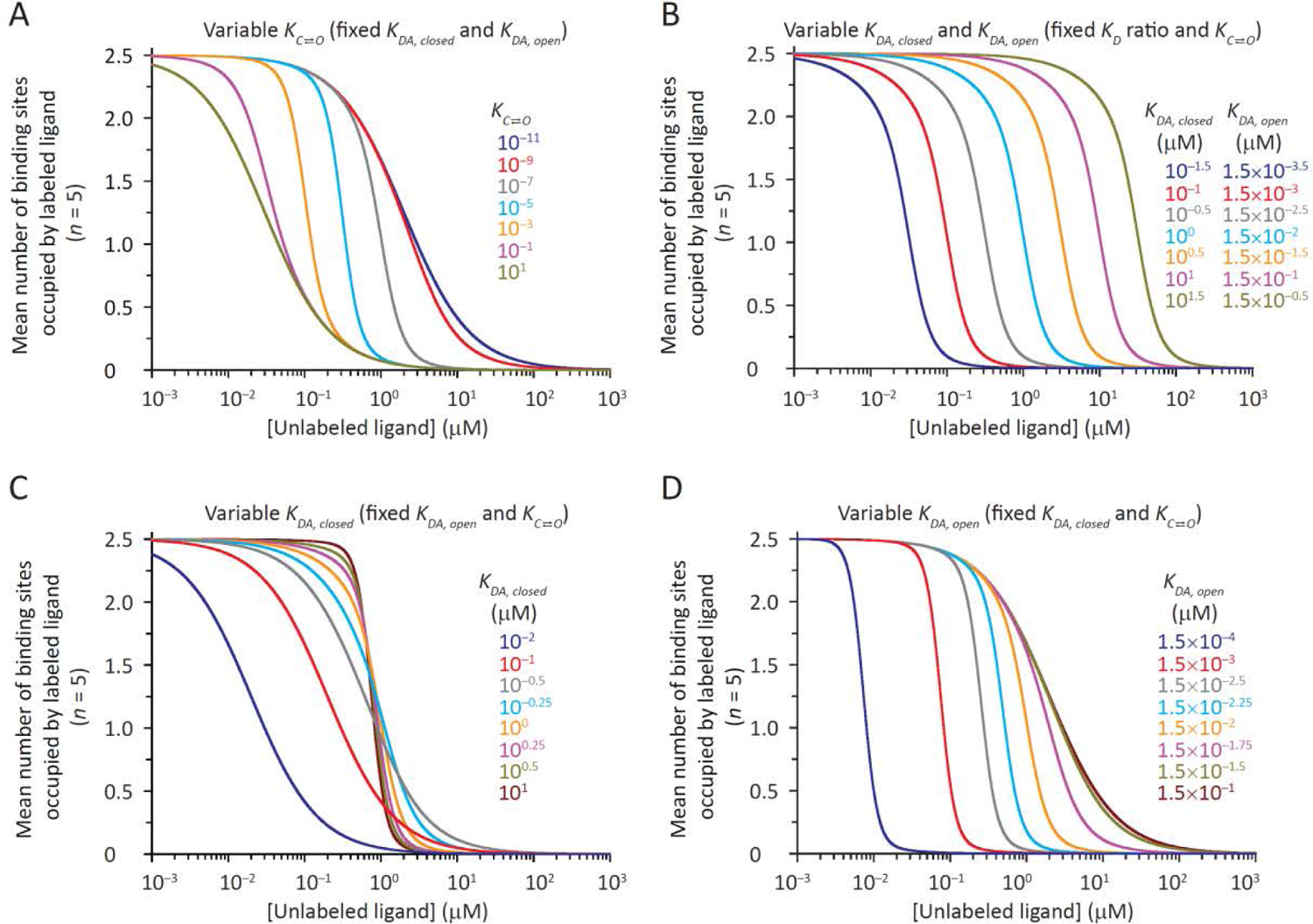
Calculated competition concentration–response curves. All curves were calculated using the reaction scheme in Figure 1. Unless otherwise stated, the parameters were: *K_C_*_⇌_*_O_* = 10^- 7^; *K_DA,closed_* = 1 μM; *K_DA,open_* = 15 nM; and *K_DB,closed_* = *K_DB,open_* = 1 nM. The ratio between the fixed and half-saturation concentrations of labeled ligand was unity, and hence, the mean number of binding sites per receptor occupied by labeled ligand at zero concentration of unlabeled ligand was one-half of their total number. (**A**–**D**) Effect of changes in the unliganded-gating equilibrium constant, the closed-state affinity, and/or the open-state affinity. In *B*, for the sake of simplicity, we assumed that the closed- and open-state affinities are linearly related (*K_DA,open_* = factor×*K_DA,closed_*) in such a way that changes in the former resulted in changes in the latter, but their ratio remained constant. For the muscle AChR, however, experiments have suggested that *K_DA,open_* = *K_DA,closed_*^factor^, instead (Nayak et al., 2019). In *C*, the variable (unlabeled-) ligand-dissociation equilibrium constant and the liganded-gating equilibrium constants change in the same direction (*Equation 6*), whereas in *D*, they do so in opposite directions. For all four panels, the concentration of unlabeled ligand (on the *x*-axes) corresponds to the concentration of unbound unlabeled ligand at equilibrium.

**Figure 5.**
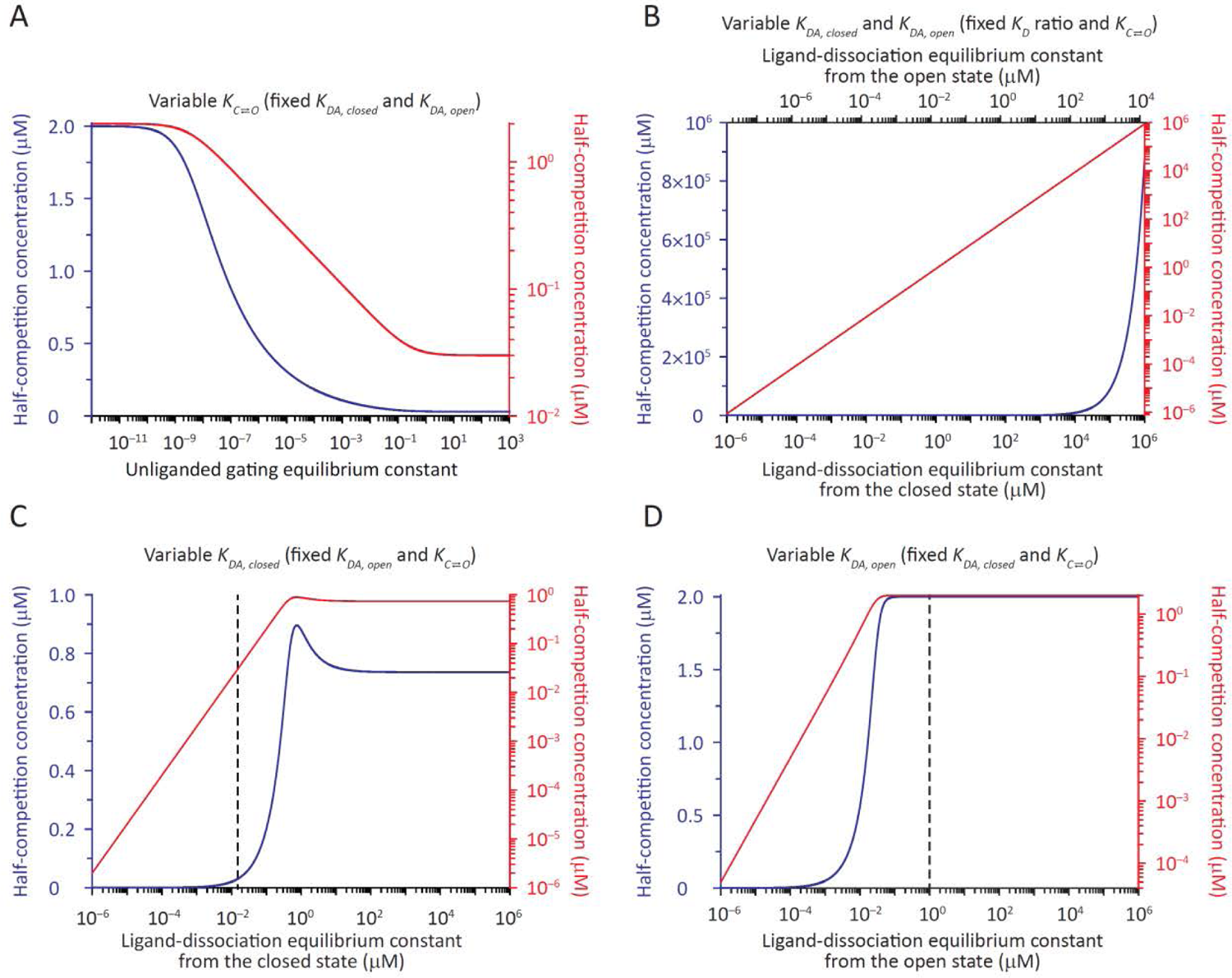
Dependence of the half-competition concentration on the unliganded-gating and (unlabeled-) ligand-dissociation equilibrium constants. All calculations were performed using the reaction scheme in Figure 1 as indicated for Figure 4. (**A**–**D**) Effect of changes in the unliganded-gating equilibrium constant, the closed-state affinity, and/or the open-state affinity. The four panels pertain to those in Figure 4. Because half-competition concentration values depend on both closed-/open-state affinities and gating, changes in this empirical parameter (upon, say, mutations) cannot be unequivocally ascribed to changes in specific equilibrium constants without additional information. The *y*-axis is displayed in both linear (in blue) and logarithmic (in red) scales. In *C* and *D*, vertical dashed lines indicate where *K_DA, closed_* = *K_DA, open_*. The labeled ligand was assumed to be an antagonist, that is, *K_DB,closed_* = *K_DB,open_*.

**Figure 6.**
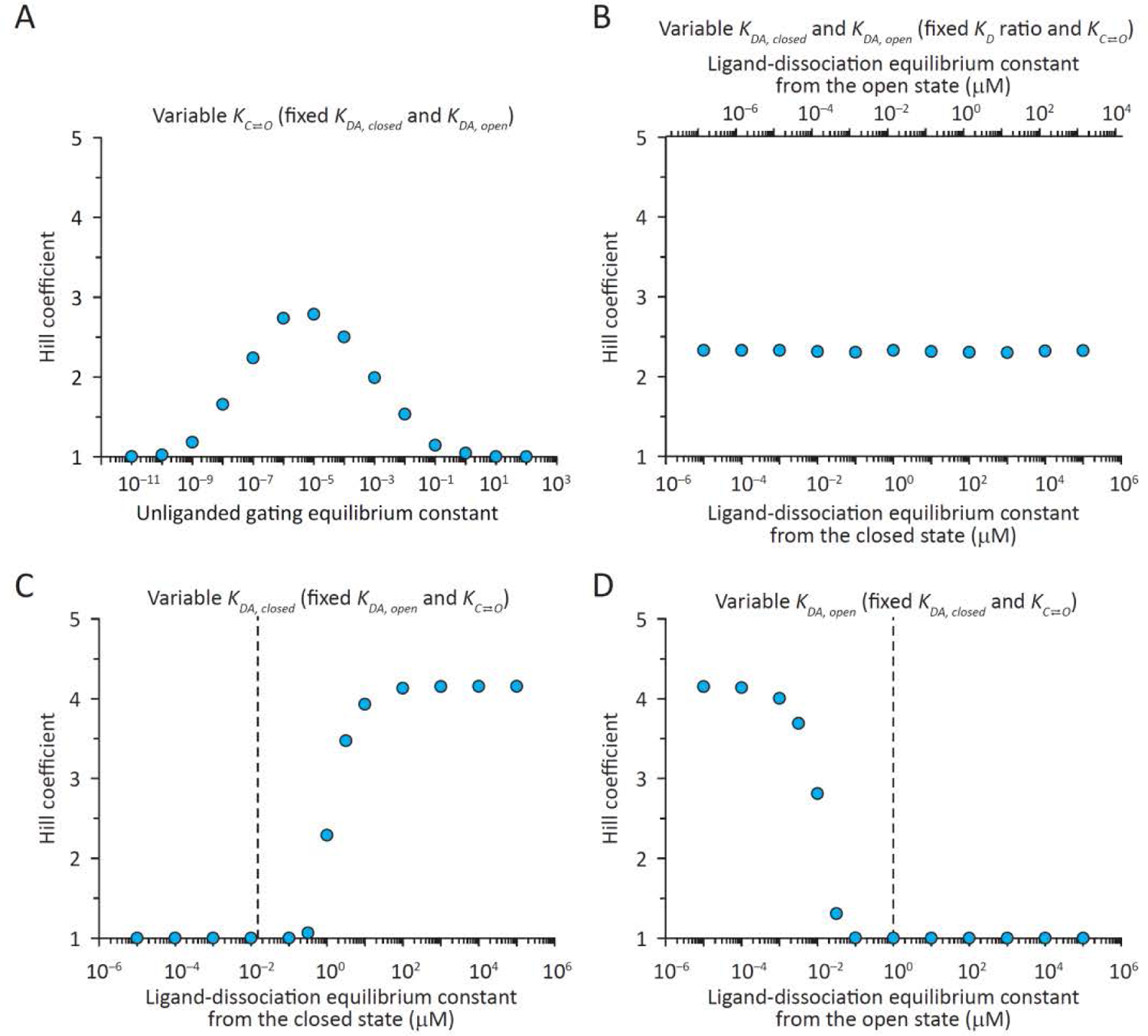
Dependence of the Hill coefficient on the unliganded-gating and (unlabeled-) ligand-dissociation equilibrium constants. (**A**–**D**) Hill-coefficient values obtained from the fitting of the calculated curves in Figure 4 with Hill equations. As expected from a receptor with identical and independent ligand-binding sites, no more than a single Hill-equation component was required to fit the equilibrium-competition curves, and the estimated values of the Hill coefficient were larger than unity and smaller than the total number of binding sites. Actually, note that the Hill coefficient remained well below 5 in all four panels. In *C* and *D*, vertical dashed lines indicate where *K_DA, closed_* = *K_DA, open_*. The labeled ligand was assumed to be an antagonist, that is, *K_DB,closed_* = *K_DB,open_*.

Figures 4 A, 5 A and 6 A show the effect of changes in the unliganded-gating equilibrium constant (without concomitant changes in ligand affinities) for the case in which the labeled ligand is an “ideal” antagonist (that is, a ligand with identical affinities for all conformations) and the unlabeled ligand, an agonist. From *Equation 6*, as the unliganded-gating equilibrium constant increases, so do the gating equilibrium constants of the receptor in its different ligation states. As a result, competition concentration–response curves shift to lower concentrations (Fig. 4 A), as expected from the higher affinity of agonists for the open-channel conformation. The displacement of these curves along the concentration axis is bounded by two limits: the half-competition concentration cannot be higher than 2×*K_D,closed_* or lower than 2×*K_D,open_* (where the *K_D_* values are those of the unlabeled ligand; Fig. 5 A). The Hill coefficient, on the other hand, approaches unity at very low and very high values of the unliganded gating equilibrium constant, going through a maximum somewhere in between (Fig. 6 A). It could be argued, however, that an inverse agonist is a better model of labeled ligand in the particular context of AChRs. Indeed, electrophysiological studies of the wild-type muscle AChR (Jackson, 1984) and gain-of-function mutants of the α7-AChR (Bertrand et al., 1997) have revealed that the binding of αBgTx reduces the spontaneous open probability (that is, it favors a non-conductive conformation), and structural models of the αBgTx-bound (Noviello et al., 2021) and unliganded α7-AChRs (Zhao et al., 2021) suggest that this non-conductive conformation is the closed (rather than the desensitized) state. Thus, we explored the behavior of competition curves for inverse-agonist labeled ligands. Figure 7 shows that, in this case, the half-competition concentration also goes from 2×*K_D,closed_* to 2×*K_D,open_*, but passing through a minimum that becomes increasingly pronounced as the inverse agonism of the labeled ligand increases. The behavior of the Hill coefficient, on the other hand, is very similar to that observed in the case of an antagonist labeled ligand, but the peak is higher and displaced to higher concentrations.

**Figure 7.**
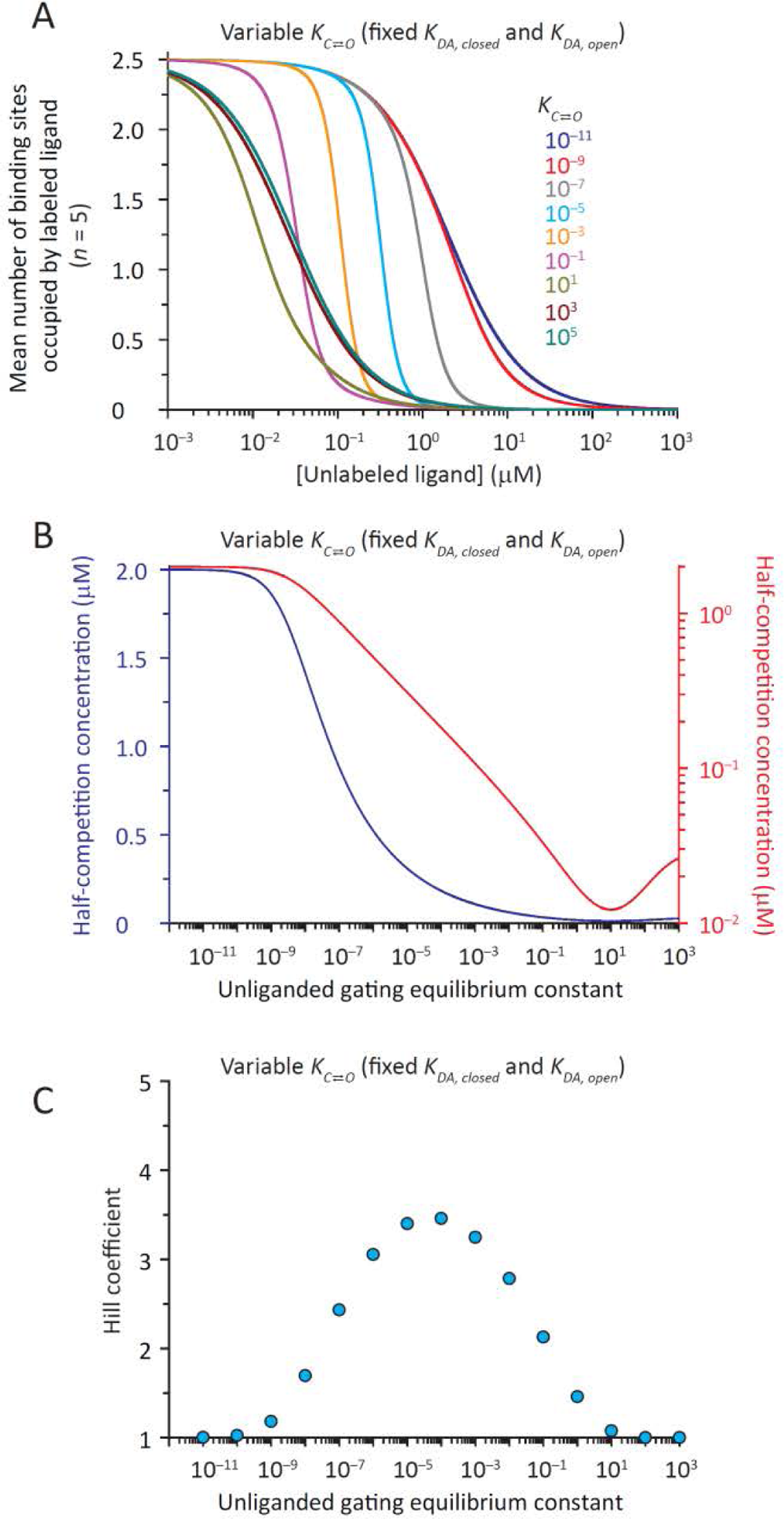
Competition concentration–response curves calculated for different values of the unliganded-gating equilibrium constant assuming the use of an inverse agonist as the labeled ligand. (**A**) Competition curves calculated using the reaction scheme in Figure 1. The parameters were: *K_DA,closed_* = 1 μM; *K_DA,open_* = 15 nM; *K_DB,closed_* = 1 nM; and *K_DB,open_* = 4 nM. The ratio between the fixed and half-saturation concentrations of labeled ligand was unity, and hence, the mean number of binding sites per receptor occupied by labeled ligand at zero concentration of unlabeled ligand was one-half of their total number. The concentration of unlabeled ligand, on the *x*-axis, corresponds to the concentration of unbound unlabeled ligand at equilibrium. (**B**) Half-competition concentration. The *y*-axis is displayed in both linear (in blue) and logarithmic (in red) scales. The plot’s minimum (most clearly displayed in the logarithmic-axis representation) becomes more pronounced as the *K_DB,open_*/*K_DB,closed_* ratio of the (inverse-agonist) labeled ligand increases. (**C**) Hill coefficient.

Figures 4 B, 5 B and 6 B show the effect of variable closed- and open-state affinities for an agonist unlabeled ligand in the idealized case when these affinities change in such a way that the ratio between them remains constant. From *Equation 6*, when this ratio and the unliganded-gating equilibrium constant remain unchanged, so do the liganded-gating equilibrium constants. Hence, as the dissociation equilibrium constants increase (that is, as the affinities decrease), competition curves shift to higher concentrations. Half-competition concentrations increase linearly as the dissociation equilibrium constants do, irrespective of whether the labeled ligand is an antagonist or an inverse agonist. The Hill coefficient, on the other hand, remains unchanged, thus showing its dependence on the channel’s gating equilibrium constants rather than ligand affinities.

Figures 4 C, 5 C and 6 C show the effect of a variable closed-state affinity for the unlabeled ligand in the idealized case when this is the only affinity that changes; Figures 4 D, 5 D and 6 D show this effect for the open-state affinity. From *Equation 6*, as *K_D,closed_* increases, so do the liganded-gating equilibrium constants, whereas the opposite relationship holds true for changes in *K_D,open_*. As shown above, as the gating equilibrium constants increase, competition curves shift to lower concentrations, whereas as *K_D_*-values increase, competition curves shift to higher concentrations. In Figure 4, C and D, and Figure 5, C and D, however, both liganded gating and closed- or open-state affinities change, and the curves shift to higher concentrations with increasing *K_D_*-values regardless of whether the liganded gating equilibrium constants concomitantly increase (Fig. 4 C and Fig. 5 C) or decrease (Fig. 4 D and Fig. 5 D); this reveals the dominant effect of ligand affinities over gating on the position of competition curves along the concentration axis. Hill-coefficient values, on the other hand (Fig. 6, C and D), change in opposite directions—increasing, in one case, and decreasing, in the other—as expected from the opposite effects of closed- and open-state affinities on the liganded-gating equilibrium constants (*Equation 6*). Furthermore, the behavior of the competition curves illustrated in Figure 4, C and D is essentially the same irrespective of whether the labeled ligand used for the calculations is an antagonist or an inverse agonist.

The interpretation of Hill-coefficient values is often linked to the concept of cooperativity of ligand binding. In all the calculated competition curves shown above, in Figure 4, the sites were assumed to be identical and independent of each other, and thus, binding-site affinities only changed as a result of the global closed ⇌ open state transition—ligand affinities did not change as a function of occupancy within a given end state of the receptor-channel. Under these particular conditions, a Hill coefficient can be thought of as a measure of the extent to which the composition of the mixture of states populated by the channel (specifically, open or desensitized *versus* closed) changes between the ends of the competition curve, in going from zero to saturating concentrations of unlabeled ligand. In one of these ends, the receptor is bound only to labeled ligand (to a degree that depends on the latter’s fixed concentration), and in the other, it is fully bound to unlabeled ligand. As a result, in these assays, Hill-coefficient values depend not only on properties of the receptor and its interaction with the unlabeled ligand, but also, on the concentration of labeled ligand and the properties of the labeled-ligand–receptor complex. This “intuitive” interpretation of the Hill coefficient nicely explains, for example, the value expected for the competition curve between two antagonists (*n_H_* = 1; *Equation 3*). Indeed, in this case, the probability of the receptor being open or desensitized (rather than closed) stays unchanged throughout the curve, whether it is the labeled or the unlabeled ligand that is bound, and thus, the coefficient takes its minimum possible number. On the other hand, changes in the open/desensitized probability are more pronounced for the competition between unlabeled agonists and labeled antagonists, and thus, for these curves, 1 < *n_H_* < *n*. Finally, these ideas also help us understand the behavior of the Hill coefficient in, for example, Figure 6 A. Here, *n_H_* approaches unity at both very low and very high values of the unliganded gating equilibrium constant, and 1 < *n_H_* < *n* otherwise. This is because, at sufficiently extreme values of this gating equilibrium constant, the channel is essentially closed or essentially open/desensitized throughout the competition curve regardless of whether the ligands are antagonists, agonists or inverse agonists. Evidently, larger-than-unity Hill-coefficient values do not necessarily imply that ligand binding-site affinities increase, within a given end state of the receptor-channel, as the occupancy of receptor sites increases.

### Practical implementation of an assay: time, temperature, and ligand depletion

Although ligand-binding assays at equilibrium have been in use for several decades now, published protocols and reported results for α-BgTx-binding AChRs span a wide range. In particular, an analysis of the literature revealed that the critical distinction between the short incubations required for protection assays and the long incubations required for equilibrium assays is often blurred, and that the Hill coefficient is frequently reported to take values that lie outside the expected bounds. Thus, we set out to identify the optimal conditions for equilibrium-binding assays.

Saturation and competition assays were performed on resuspended HEK-293 cells transiently expressing the wild-type human α7-AChR or some mutants thereof. The labeled ligand was [^125^I]-α-BgTx, and its fixed (unbound) concentration in competition assays was chosen to be equal to its half-saturation concentration, which in turn, was estimated from saturation-binding assays (Fig. 8). After an incubation period, the reactions were terminated by centrifugation, which separated cell-bound label from unbound, “free” label. To reduce non-specifically bound toxin, the cell pellets were washed extensively with a sodium-saline solution, a crucial step made possible by the toxin’s slow dissociation from the α7-AChR. Inconveniently, however, the slow dissociation kinetics of α-BgTx also slowed down the approach to equilibrium.

**Figure 8.**
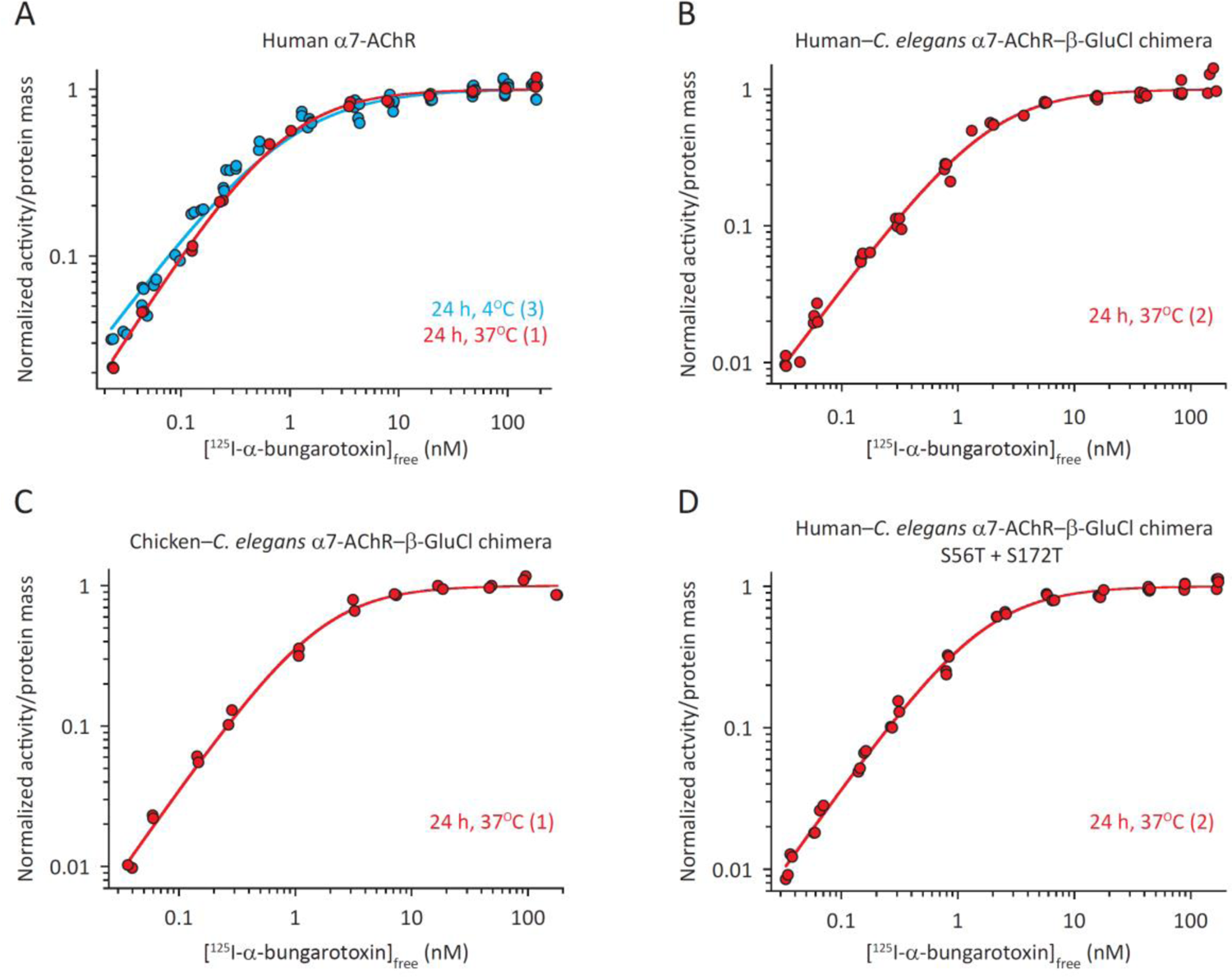
Experimental saturation concentration–response curves for the human α7-AChR and related constructs. (**A**) Human α7-AChR. (**B**) Human–*C. elegans* α7-AChR–β-GluCl ECD–TMD chimera. (**C**) Chicken–*C. elegans* α7-AChR–β-GluCl ECD–TMD chimera. (**D**) Human–*C. elegans* α7-AChR–β-GluCl chimera S56T + S172T. All curves were best fitted with one-component Hill equations. Moreover, for the four curves at 37°C, *n_H_* ≅ 1, as expected at equilibrium from an inverse agonist binding to a receptor that is essentially closed when unliganded and has identical and independent ligand-binding sites. The number of independent saturation assays contributing to each plotted curve is indicated, in parentheses, in the corresponding figure caption. For all four panels, the normalized radioactivity (on the *y*-axes) represents specifically bound toxin calculated by subtracting the amount of non-specifically bound toxin from the total cell pellet-associated radioactivity. The concentration of [^125^I]-α-BgTx (on the *x*-axes) corresponds to the concentration of unbound toxin, which was estimated from the radioactivity measured in the supernatant of each individual binding reaction. Half-saturation and Hill-coefficient values estimated from assays incubated at 37°C for 24 h are shown in Table 1.

**Table 1.**
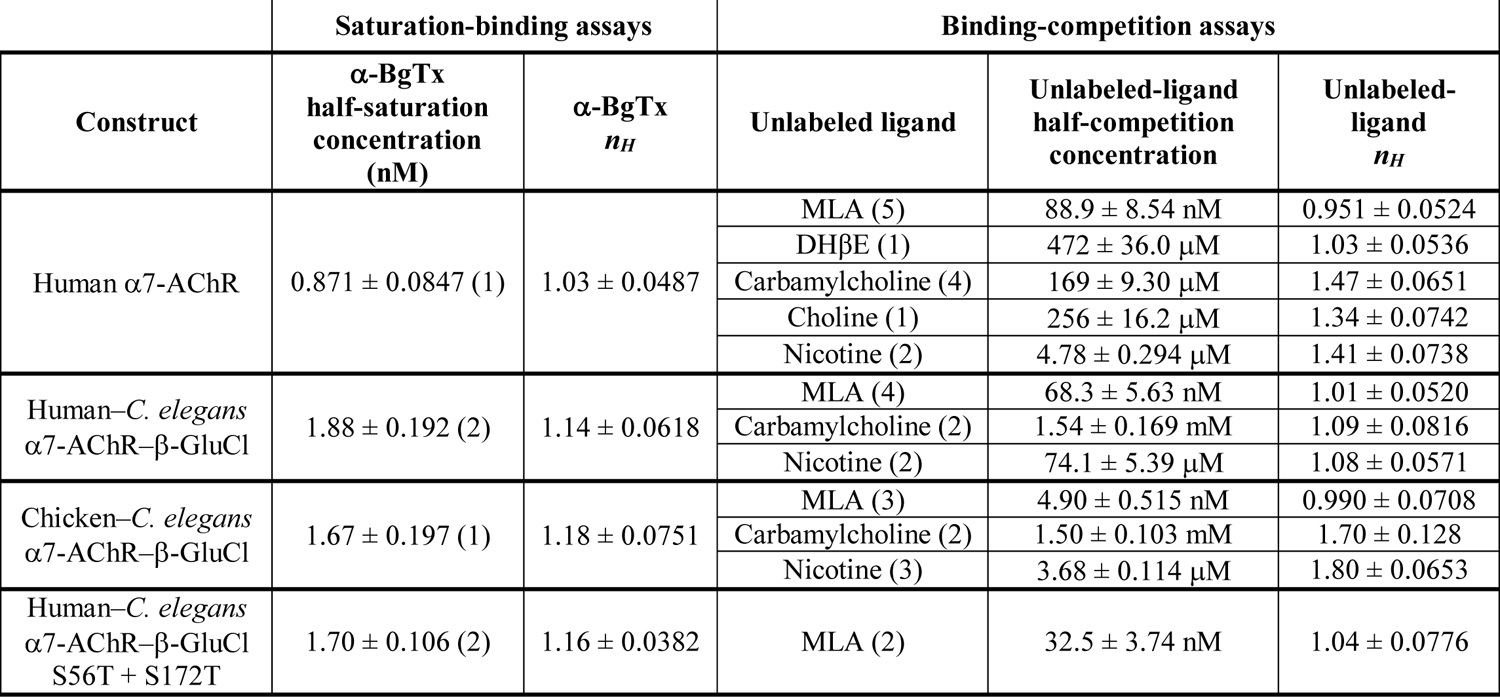
Half-effective concentrations and Hill-coefficient values for various cholinergic ligands and constructs of the human α7-AChR obtained from incubations at 37°C Saturation-binding reactions were incubated at 37°C for 24 h, whereas binding-competition reactions were incubated at 37°C for 24 or 48 h. All individual curves for a given combination of construct and ligand were globally fitted (regardless of incubation duration), and standard errors were obtained from these fits. For competition experiments, the ratio between the fixed and half-saturation concentrations of [^125^I]-α-BgTx was approximately unity. Parameter estimates (mean ± 1 SE) are presented with three significant figures. The number of independent saturation or competition assays contributing to each parameter estimation is indicated in parentheses. Parameter estimates obtained from incubations at 4°C or 22°C are not listed.

Figure 9 shows the effects of duration and temperature of the incubations on the competition between α-BgTx and methyllycaconitine (MLA; an inverse agonist; (Bertrand et al., 1997)) or the non-hydrolyzable, synthetic ACh analog carbamylcholine (an agonist), whereas Figure 10 shows the effect of temperature on the competition between α-BgTx and choline (an agonist), nicotine (an agonist) or dihydro-β-erythroidine (DHβE; an extremely weak agonist; (Bertrand et al., 1997)). For all five competing ligands (whose structures are shown in Fig. 11), the effects were qualitatively the same: as the temperature rose from 4°C to 37°C, and the duration increased from 4 h to 48–96 h, the Hill equations that best fitted the data switched from having two components to having only one, and the curves seemed to shift to the right. At 4°C, 24-h incubations were too short for equilibrium to be attained, but at 37°C, 24-h incubations seemed long enough. With these results in mind, we decided to adopt an incubation temperature of 37°C and a duration of 24–48 h for these assays. Also, we found that the binding-competition curves were more sensitive to the temperature of the incubations than were the saturation-binding curves (Fig. 8 A).

**Figure 9.**
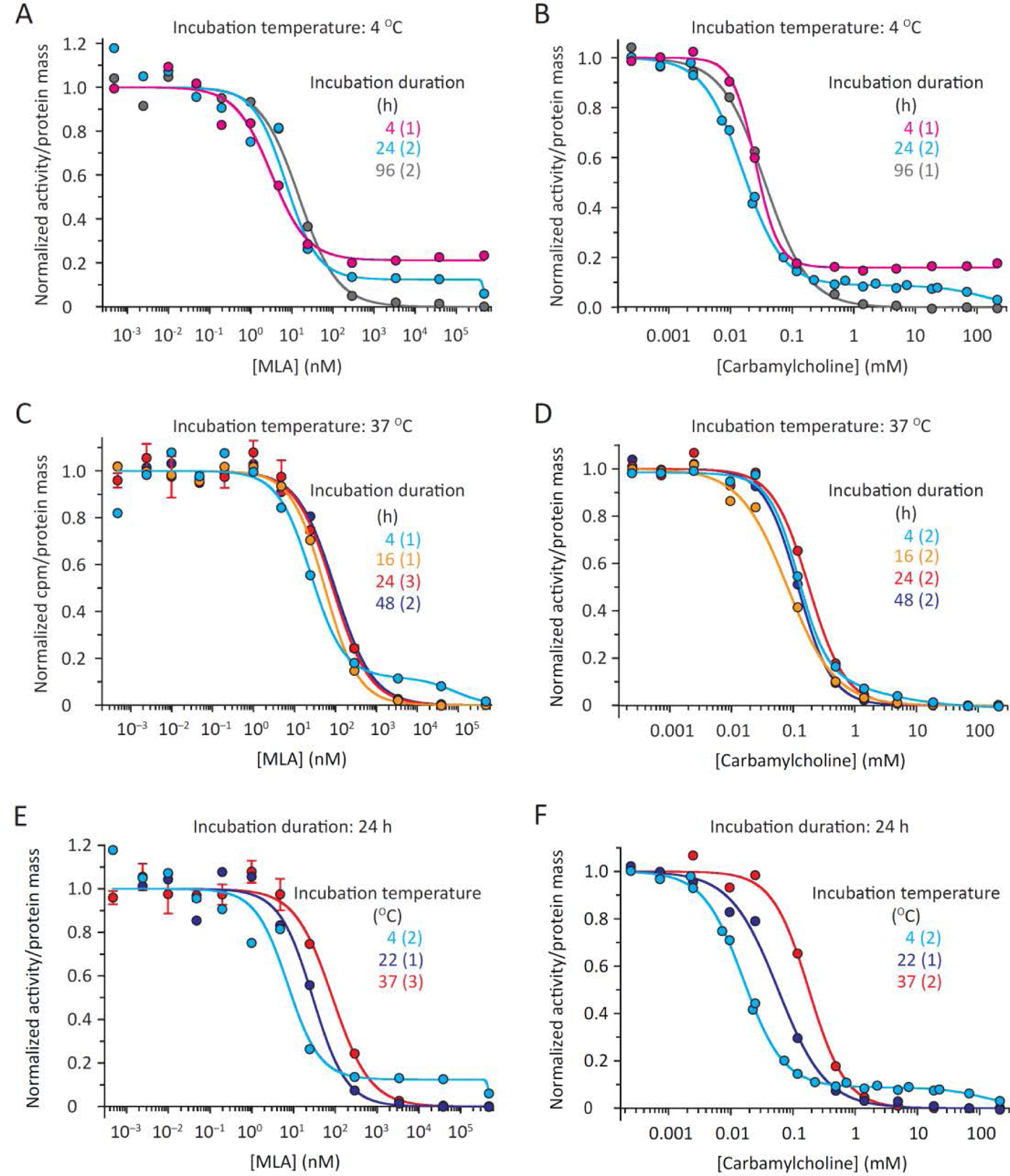
Time and temperature dependence of experimental competition concentration– response curves for MLA and carbamylcholine on the human α7-AChR. (**A**, **B**) 4°C, variable incubation times. MLA is an inverse agonist, whereas carbamylcholine is an agonist. Only the curves corresponding to 96-h incubations were best fitted with one-component Hill equations; all others required a second component. (**C**, **D**) 37°C, variable incubation times. The curves corresponding to 4-h incubations were best-fitted with two-component Hill equations; all others, with one. (**E**, **F**) 24-h, variable incubation temperatures. The (re-plotted) curves at 4°C and 37°C are those in *A*–*D*. For all assays, the ratio between the fixed and half-saturation concentrations of [^125^I]-α-BgTx was approximately unity. The number of independent competition assays contributing to each plotted curve is indicated, in parentheses, in the corresponding figure caption; errors were calculated only when the latter was larger than 2. Error bars (± 1 SE) smaller than the size of the symbols were omitted. For all six panels, the concentration of unlabeled ligand (on the *x*-axes) corresponds to the total (bound plus unbound) concentration. Under the conditions of our experiments, the depletion of unlabeled ligand owing to its binding to the receptor was inferred to be low, and thus, the total concentration was deemed to be a good approximation for the concentration of unbound unlabeled ligand at equilibrium. Half-competition and Hill-coefficient values estimated from assays incubated at 37°C for 24 or 48 h are shown in Table 1.

**Figure 10.**
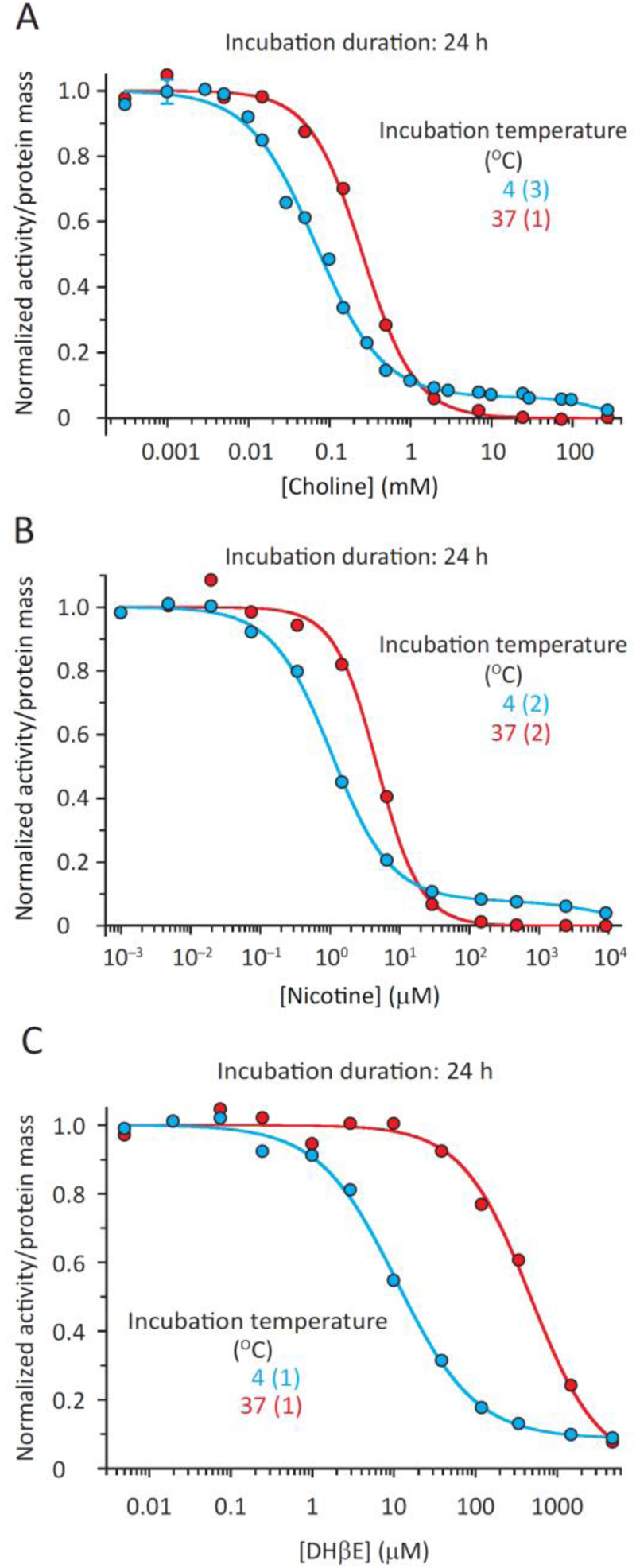
Temperature dependence of experimental competition concentration–response curves for choline, nicotine, and DHβE on the human α7-AChR. (**A**) Choline, an agonist. (**B**) Nicotine, an agonist. (**C**) DHβE, an extremely weak agonist. The three curves at 4°C were best fitted with two-component Hill equations, whereas the three curves at 37°C, with one-component Hill equations. For all assays, the ratio between the fixed and half-saturation concentrations of [^125^I]-α-BgTx was approximately unity. The number of independent competition assays contributing to each plotted curve is indicated, in parentheses, in the corresponding figure caption; errors were calculated only when the latter was larger than 2. Error bars (± 1 SE) smaller than the size of the symbols were omitted. For all three panels, the concentration of unlabeled ligand (on the *x*-axes) corresponds to the total (bound plus unbound) concentration. Under the low ligand-depletion conditions of our experiments, this concentration was deemed to be a good approximation for the concentration of unbound unlabeled ligand at equilibrium. Half-competition and Hill-coefficient values estimated from assays incubated at 37°C for 24 h are shown in Table 1.

**Figure 11.**
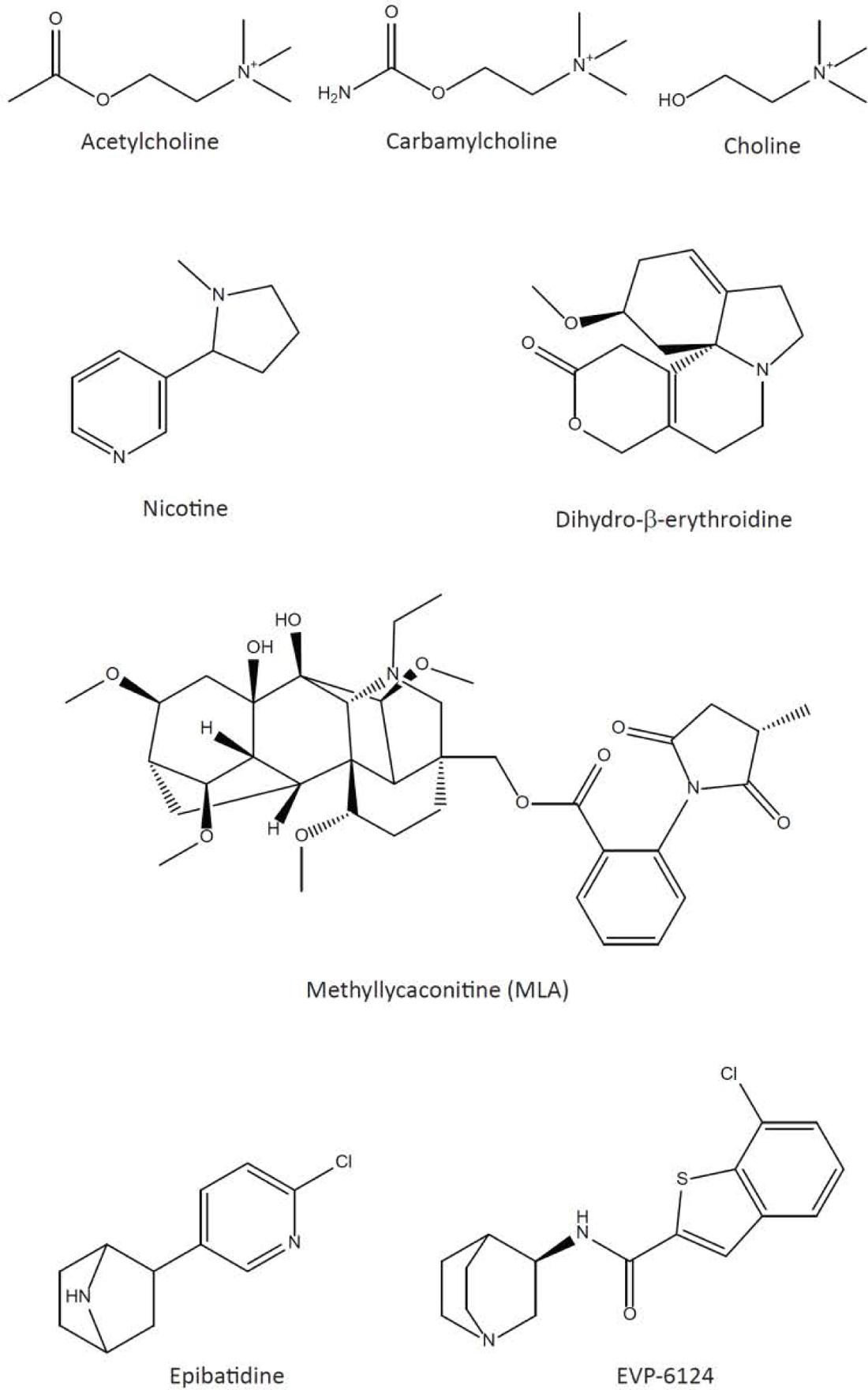
Structures of orthosteric ligands used or discussed in this work.

Needless to say, in addition to the need of measuring the amount of bound signal, the concentrations of unbound labeled and unlabeled ligands at equilibrium need to be known for all the individual points of the curve. Indeed, the concentration of unbound labeled ligand needs to remain constant, whereas the concentration of unbound unlabeled ligand is the independent variable plotted on the *x*-axis. Although the concentration of unbound labeled ligand can be easily measured upon separating it from the bound form (in our case, in the supernatant; Fig. 12), measuring the concentration of unbound unlabeled ligand is often more cumbersome, and thus, we assumed its value to be equal to its total concentration (total = bound + unbound). Clearly, for the latter assumption to be valid, the concentration of receptors needs to be low enough to make the depletion of ligand negligible. This is a challenge because, at the same time, the amount of receptors needs to be large enough to generate a sizable bound signal. We addressed this issue by trial and error—adjusting the amount of cDNA used in the transfections and the number of transfected cells—so as to strike a balance. We deemed the concentration of ligand-binding receptor sites to be adequate when: a) The difference between the maximum and minimum values of the measured concentrations of unbound labeled toxin for any given competition curve (Fig. 12) was <0.2 nM, and the mean was lower than the total concentration of added toxin by <0.2 nM (for the α7-AChR and its mutants, the desired concentration of unbound α-BgTx was in the ∼1–2 nM range); and b) The specific cell-bound radioactivity in the absence of unlabeled ligand was higher than the non-specifically bound radioactivity by a factor >10. In our assays, these conditions were met when the concentration of toxin-binding sites was in the 0.03–0.3 nM range. Owing to their different expression levels, different constructs required different conditions to hit these values (see Materials and methods), but once identified, they remained reproducibly valid for all subsequent assays.

**Figure 12.**
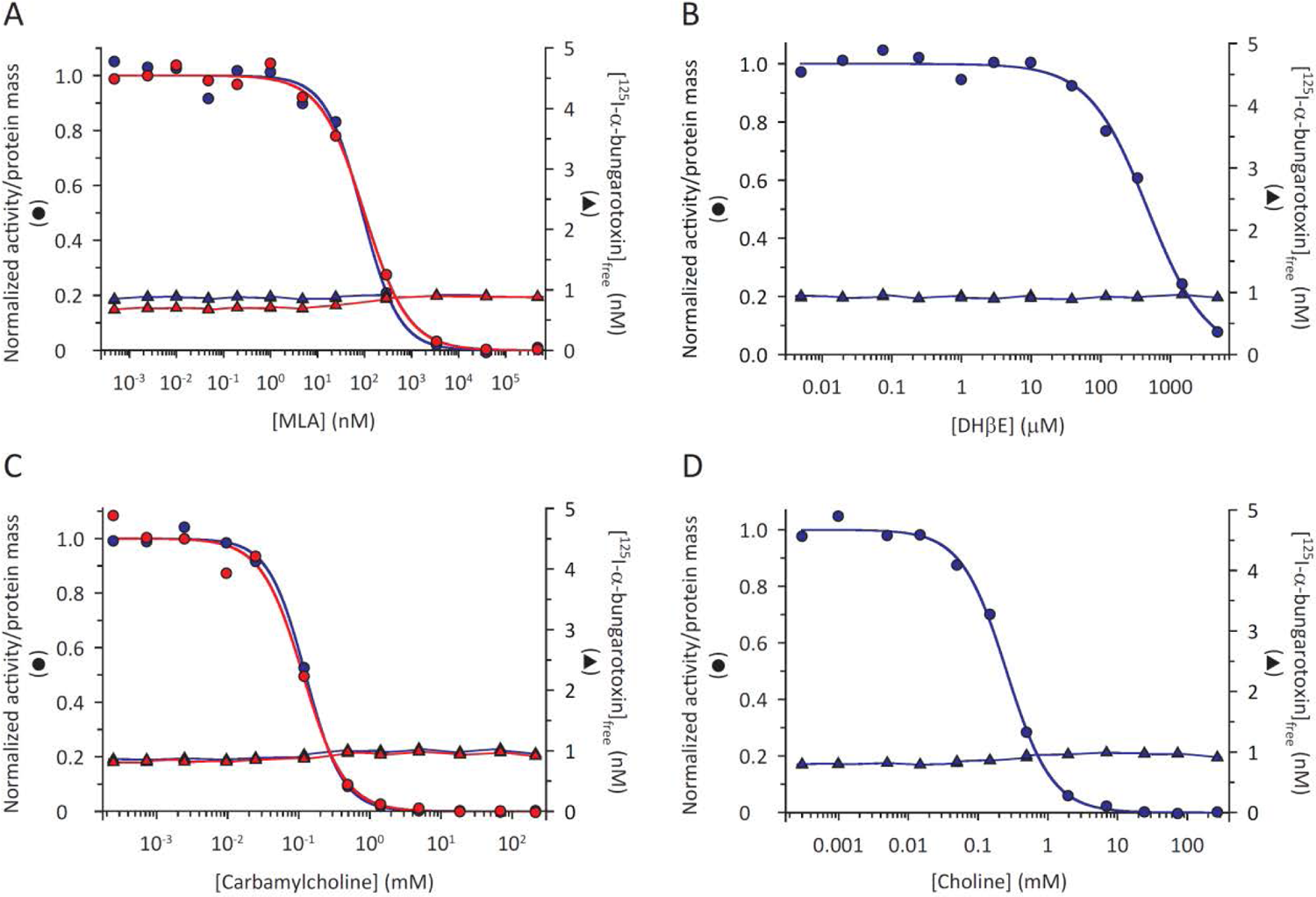
Experimental competition concentration–response curves from individual assays and the concentration of unbound [^125^I]-α-BgTx. The receptor was the wild-type human α7-AChR. (**A**) MLA. (**B**) DHβE. (**C**) Carbamylcholine. (**D**) Choline. In *A* and *C*, different colors represent different individual assays. Upon completion of the incubation period (here, 24 h at 37°C for all four panels), cell-bound [^125^I]-α-BgTx was physically separated from unbound [^125^I]-α-BgTx by centrifugation. The (normalized) radioactivity associated with the former (circles) was plotted on the left *y*-axes, whereas that associated with the latter (triangles), expressed in concentration units, was plotted on the right *y*-axes. For all four panels, the ratio between the fixed and half-saturation concentrations of [^125^I]-α-BgTx in each assay was approximately unity. The concentration of unlabeled ligand (on the *x*-axes) corresponds to the total (bound plus unbound) concentration. Under the low ligand-depletion conditions of our experiments, this concentration was deemed to be a good approximation for the concentration of unbound unlabeled ligand at equilibrium.

At 37°C and with 24-h incubations, the α-BgTx saturation curve of the human α7-AChR in intact HEK-293 cells was best fitted with a one-component Hill equation with *n_H_* = 1.03 ± 0.05 and a half-saturation concentration of 0.87 ± 0.09 nM (Fig. 8 and Table 1). The value of the Hill coefficient is consistent with the toxin’s inverse agonism on a receptor that displays a nearly undetectable unliganded channel activity and whose closed-state toxin-binding sites are identical and independent of each other’s occupancy. Moreover, under these particular conditions, *Equation 5* provides an excellent description of the ligand–receptor interaction, and thus, the toxin’s half-saturation concentration is a good estimate of its *K_D_* from the closed state. Our estimate of the latter’s value agrees most closely with those of Oz and coworkers (1.18 ± 0.29 nM; (Shabbir et al., 2021)), Sullivan and coworkers (0.71 ± 0.11 nM; (Gopalakrishnan et al., 1995)), and Lindstrom and coworkers (0.81 nM; (Peng et al., 1994)) for the same receptor in intact SH-EP1 cells, HEK-293 cell-membrane homogenates, and detergent micelles, respectively.

The competition between α-BgTx and MLA or DHβE for binding to the wild-type α7-AChR (at 37°C, 24 or 48 h) gave rise to concentration–response curves that are best fit with a single Hill-equation component of *n_H_* ≅ 1 (Fig. 9, C and E; Fig. 10 C; and Table 1). This is further experimental evidence for the notion that the five closed-state ligand-binding sites (the open state is hardly visited in competitions between these ligands) have indistinguishable affinities and are independent of each other’s occupancy. Certainly, even a small degree of positive cooperativity among sites—only strong enough to make the affinity of the tetra-liganded channel for the fifth molecule of antagonist/inverse-agonist ligand appreciably higher—would be expected to increase the Hill coefficient above unity (Fig. 13). Similarly, even a small degree of negative cooperativity would have been detected as a competition curve that requires a lower-than-unity Hill coefficient, or even a second Hill-equation component, to be best fitted (Fig. 13).

**Figure 13.**
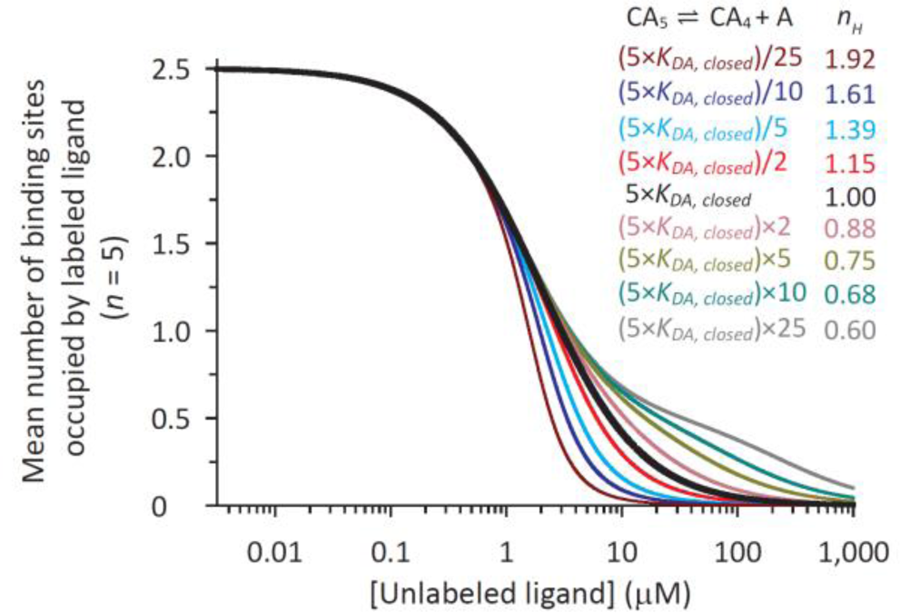
Effect of positive or negative cooperativity of binding on the competition between a labeled inverse agonist and an unlabeled antagonist. Curves were calculated using the reaction scheme in Figure 1 assuming independence of sites (in black) or the occurrence of different degrees of deviations from it (“cooperativity”). Cooperativity was assumed to be caused by, and to only affect, the binding of unlabeled ligand; binding of the labeled ligand, on the other hand, was assumed to neither cause nor be affected by these departures from independence. Furthermore, cooperativity was assumed to be weak; it was only strong enough to make the affinity of the tetra-liganded channel for the fifth molecule of antagonist appreciably different from 5×*K_DA,closed_*, that is, the value expected from independence of five identical sites. Also, for the sake of simplicity, cooperativity was assumed to affect the affinities of the closed and open states by the same factor. Unless otherwise stated, the parameters used for these calculations were: *K_C_*_⇌_*_O_* = 10^-7^; *K_DA,closed_* = *K_DA,open_* = 1 μM; *K_DB,closed_* = 1 nM; and *K_DB,open_* = 4 nM. The ratio between the fixed and half-saturation concentrations of labeled ligand was unity, and hence, the mean number of binding sites per receptor occupied by labeled ligand at zero concentration of unlabeled ligand was one-half of their total number. The extent to which the affinity of the receptor bound to four molecules of unlabeled ligand for a fifth molecule of unlabeled ligand differs from the value expected from independence is indicated for each curve as the dissociation equilibrium constant of the fully bound, pentaliganded channel. The Hill-coefficient values obtained from the fits of the calculated curves with one-component Hill equations are also indicated. The concentration of unlabeled ligand, on the *x*-axis, corresponds to the concentration of unbound unlabeled ligand at equilibrium.

Conversely, as elaborated in the sections above, the larger-than-unity Hill-coefficient values required to fit the competition curves between α-BgTx and carbamylcholine, choline or nicotine (at 37°C, 24 or 48 h; Fig. 9, D and F; Fig. 10, A and B; and Table 1) do not necessarily imply the occurrence of positive cooperativity among sites; instead, they may simply reflect the fact that these unlabeled ligands are agonists.

When binding reactions are not allowed to reach equilibrium, competition curves often display features that are theoretically inconsistent with the notion of identical and independent binding sites. Indeed, out-of-equilibrium concentration–response curves often require two (or more) Hill-equation components or one component with *n_H_* < 1 to be best fitted (Fig. 9; Fig. 10; and Fig. 14). Quite notably, these theoretically nonsensical features would be expected at equilibrium from ligand-binding proteins whose binding-site affinities display various degrees of interdependence in the form of “negative cooperativity” (Fig. 15), and therefore, could be misinterpreted as genuine signs of interactions between sites. For proteins with identical and independent binding sites, however, these anomalous features only indicate that the binding reactions were terminated too soon.

**Figure 14.**
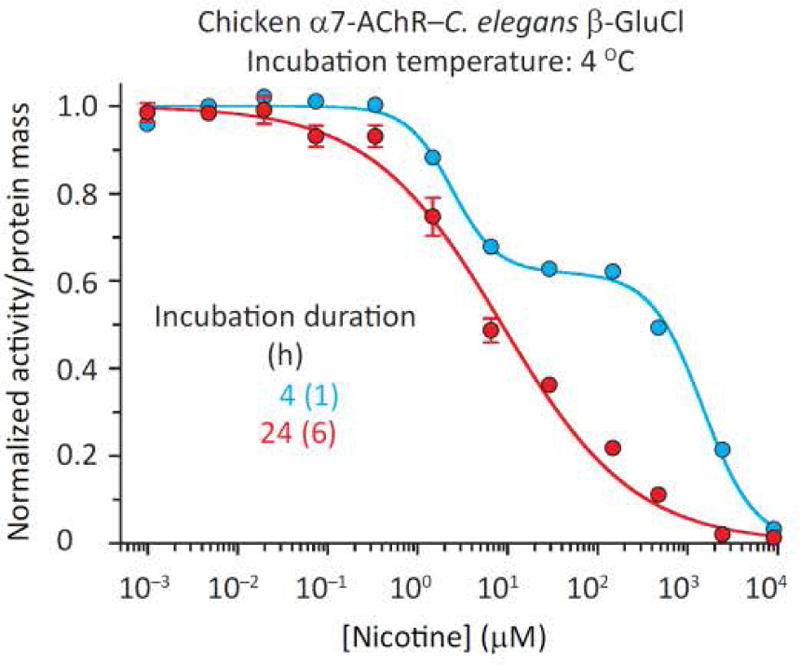
Anomalous features in experimental α-BgTx–agonist-competition curves. The receptor was the chicken–*C. elegans* α7-AChR–β-GluCl chimera. The ratio between the fixed and half-saturation concentrations of [^125^I]-α-BgTx in each assay was approximately unity. The curve in cyan was best fitted with a two-component Hill equation, whereas that in red, with a one-component Hill equation where *n_H_* = 0.59. The number of independent competition assays contributing to each plotted curve is indicated, in parentheses, in the corresponding figure caption; errors were calculated only when the latter was larger than 2. Error bars (± 1 SE) smaller than the size of the symbols were omitted. The concentration of (unlabeled) nicotine, on the *x*-axis, corresponds to the total (bound plus unbound) concentration. Under the low ligand-depletion conditions of our experiments, this concentration was deemed to be a good approximation for the concentration of unbound nicotine at equilibrium.

**Figure 15.**
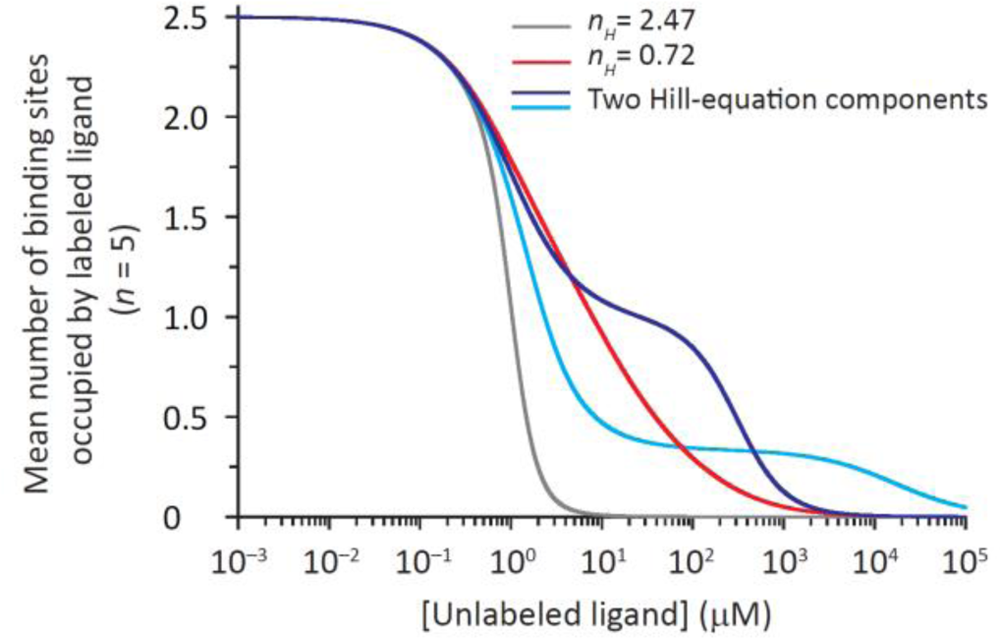
Effect of negative cooperativity of binding on the competition between a labeled inverse agonist and an unlabeled agonist. The curves were calculated using the reaction scheme in Figure 1 assuming independence of sites (in gray) or the occurrence of different degrees of deviations from it (“cooperativity”). Negative cooperativity was assumed to be caused by, and to only affect, the binding of unlabeled ligand; binding of the labeled ligand, on the other hand, was assumed to neither cause nor be affected by these departures from independence. Also, for the sake of simplicity, cooperativity was assumed to affect the affinities of the closed and open states by the same factor. Unless otherwise stated, the parameters used for these calculations were: *K_C_*_⇌_*_O_* = 10^-7^; *K_DA,closed_* = 1 μM; *K_DA,open_* = 15 nM; *K_DB,closed_* = 1 nM; and *K_DB,open_* = 4 nM. The ratio between the fixed and half-saturation concentrations of labeled ligand was unity, and hence, the mean number of binding sites per receptor occupied by labeled ligand at zero concentration of unlabeled ligand was one-half of their total number. The curve in cyan assumes that the affinity of the receptor bound to four molecules of unlabeled ligand for a fifth molecule of unlabeled ligand is lower than that expected from independence by a factor of 10^5^. The curve in blue assumes that the affinity of the receptor bound to four molecules of unlabeled ligand for a fifth molecule of unlabeled ligand is lower than that expected from independence by a factor of 10^3^, whereas the affinity of the receptor bound to three molecules of unlabeled ligand (whether bound to one molecule of labeled ligand or not) for a fourth molecule of unlabeled ligand is lower by a factor of 350. The curve in red assumes that the unlabeled-ligand affinity of the receptor bound to four molecules of unlabeled ligand is lower than that expected from independence by a factor of 10^3^, that of the receptor bound to three molecules of unlabeled ligand is lower by a factor of 10, and that of the receptor bound to two molecules of unlabeled ligand is lower by a factor of 2. For the curves in cyan and blue, negative cooperativity of binding is manifest as a clear second Hill-equation component, whereas for the curve in red, negative cooperativity is manifest as a shallow, single Hill-equation component best fitted with *n_H_* < 1. The curve in gray was also best fitted with a single Hill-equation component, but *n_H_* > 1. The concentration of unlabeled ligand, on the *x*-axis, corresponds to the concentration of unbound unlabeled ligand at equilibrium.

### Answering specific mechanistic questions with equilibrium-binding assays

A long-standing question in the field of NGICs is whether mutations to the TMD can affect the ligand affinities of the rather distant neurotransmitter-binding sites, in the ECD (the distance between bound orthosteric ligands and the center of the ion-channel pore is ∼50 Å). This is an example of the more general question as to how far the structural perturbations caused by side-chain mutations can travel through a protein. In the context of the muscle AChR and its naturally occurring agonist, ACh, efforts to tackle this issue with the kinetic analysis of single-channel recordings have led to diametrically opposed answers (Hatton et al., 2003; Purohit et al., 2015; Wang et al., 1997).

Here, to eliminate the complications associated with ligand affinities that change upon opening and desensitization, we estimated the closed-state *K_D_* values of the wild-type α7-AChR and some mutants for the inverse agonist MLA. Furthermore, to avoid the use of such an indirect approach to the estimation of ligand affinities as the kinetic modeling of single-channel open and shut dwell times, we performed binding-competition assays against [^125^I]-α-BgTx following the procedures and concepts elaborated above. In addition to the wild-type human α7-AChR, we studied two other constructs: a) A chimera that combines the human α7-AChR’s ECD with the TMD of the β subunit of the glutamate-gated Cl^-^ channel (β-GluCl) from *C. elegans*, as an extreme example of a human α7-AChR bearing extensive mutations in the TMD; and b) A chimera that combines the chicken α7-AChR’s ECD with the TMD of *C. elegans* β-GluCl, as an example of a chimera with multiple mutations in the ECD. More precisely, the human and chicken α7-AChR ECDs differ at 13 positions (see Materials and methods), none of which approaches the agonists epibatidine or EVP-6124 (Fig. 11) closer than 3.0 Å in existing atomic models of the ligand-bound receptor (Noviello et al., 2021; Zhao et al., 2021) (Fig. 16). As for the TMDs, those of the human α7-AChR and *C. elegans* β-GluCl are identical at only ∼40 positions out of a total of 273 (∼15%) in the α7-AChR and 190 in β-GluCl (∼21%).

**Figure 16.**
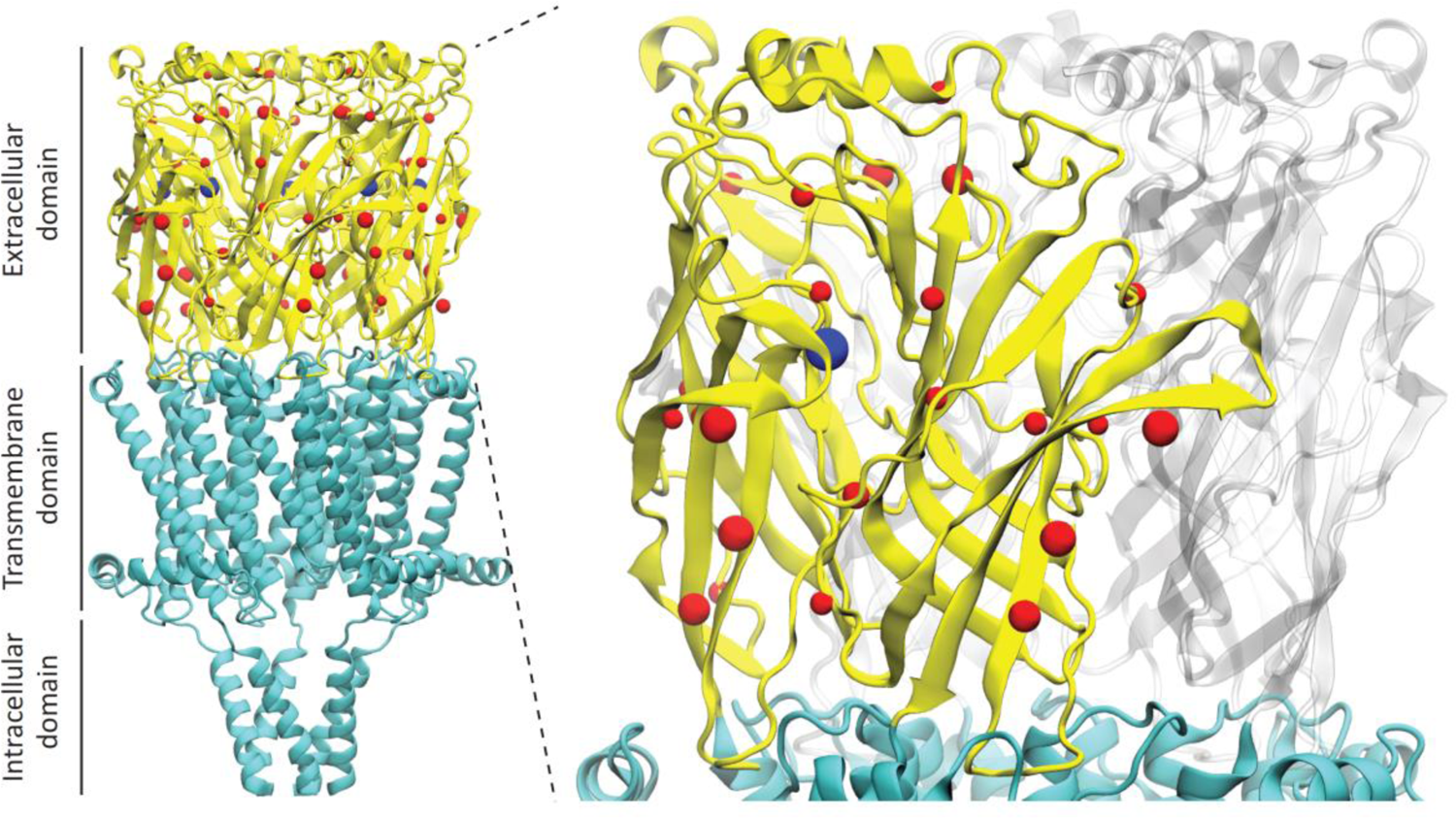
Domain architecture of pLGICs, and amino-acid differences between the human and chicken α7-AChR’s ECDs. Structural model of the human α7-AChR bound to the orthosteric agonist epibatidine and the positive allosteric modulator PNU-120596 (PDB ID: 7KOX; (Noviello et al., 2021)) displayed in ribbon representation. Blue spheres indicate the positions occupied by the five copies of bound orthosteric ligand. Red spheres indicate the location of the 13 amino-acid residues that differ between the sequences of the human and chicken α7-AChR subunits at the level of the ECD. Inset: the ECDs of two adjacent subunits, and the position occupied by the orthosteric ligand, at their interface, are emphasized. The molecular images were prepared with visual molecular dynamics (Humphrey et al., 1996).

Toxin-saturation curves for the three constructs displayed similar α-BgTx half-saturation concentrations (∼ 1–2 nM) and Hill-coefficient values (∼1; Fig. 8 and Table 1). Also, the MLA– toxin competition curves for the two constructs having a human α7-AChR ECD and highly divergent TMDs were very similar, whereas those for the two constructs having the same β-GluCl TMD and slightly different α7-AChR ECDs were clearly different (Fig. 17 A and Table 1). Under the conditions of these assays—that is, receptor-channels that barely open when unliganded, unlabeled and labeled ligands that favor the closed state, and a ratio between the fixed and half-saturation concentrations of labeled ligand equal to 1—*Equation 4* provides an excellent description of the ligand–receptor interaction. Thus, the half-competition concentration values are approximately equal to 2×*K_D,closed_*, that is, direct estimates of (true) MLA affinities.

**Figure 17.**
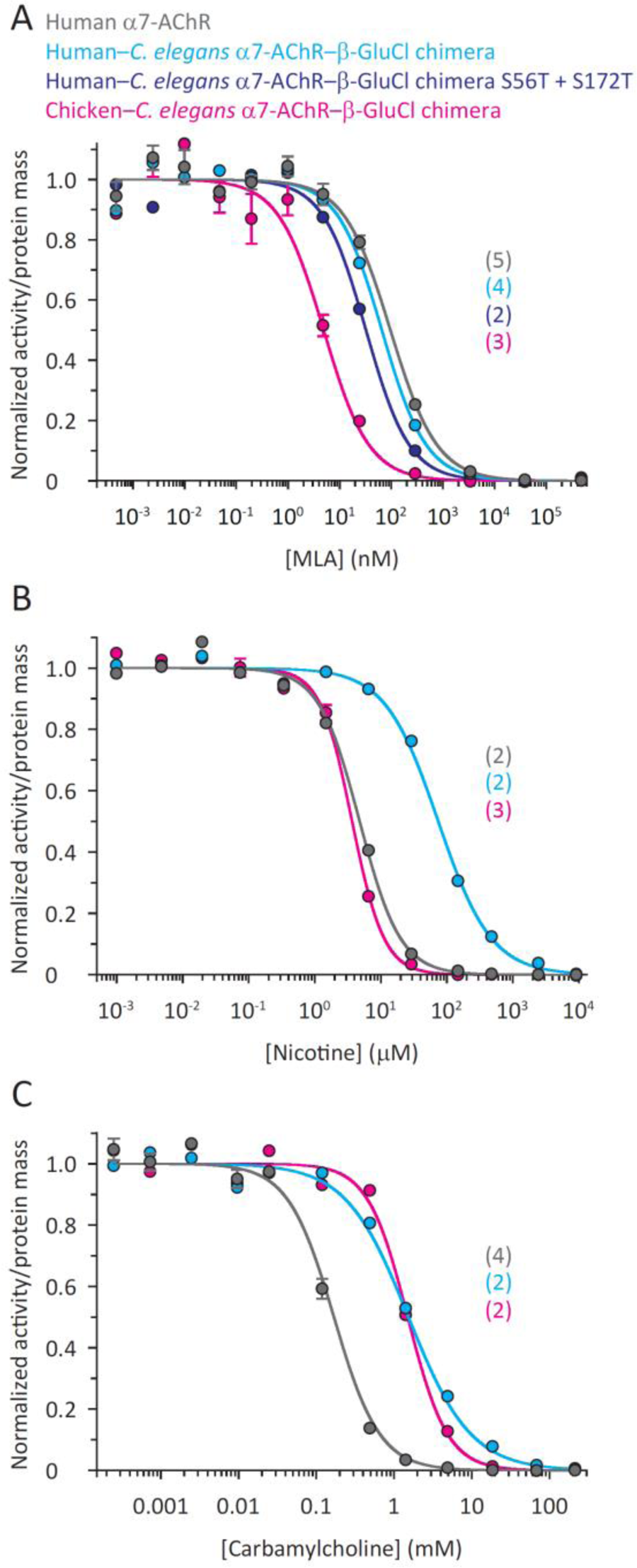
Effect of mutations to the human α7-AChR’s ECD or TMD domains on experimental competition curves. (**A**) MLA. (**B**) Nicotine. (**C**) Carbamylcholine. For all curves, the binding reactions were incubated at 37°C for 24 or 48 h. All curves were best fitted with one-component Hill equations. The ratio between the fixed and half-saturation concentrations of [^125^I]-α-BgTx in each assay was approximately unity. The number of independent competition assays contributing to each plotted curve is indicated, in parentheses, in the corresponding figure caption; errors were calculated only when the latter was larger than 2. Error bars (± 1 SE) smaller than the size of the symbols were omitted. For all three panels, the concentration of unlabeled ligand (on the *x*-axes) corresponds to the total (bound plus unbound) concentration. Under the low ligand-depletion conditions of our experiments, this concentration was deemed to be a good approximation for the concentration of unbound unlabeled ligand at equilibrium. Half-competition and Hill-coefficient values are shown in Table 1. The color code is the same for all panels. The effect of the S56T and S172T ECD mutations on the human–*C. elegans* α7-AChR–β-GluCl chimera was tested only for MLA.

Therefore, the obtained curves support the idea that small structural perturbations to the TMD do not reach the neurotransmitter binding sites (because large perturbations barely have an effect), whereas small structural perturbations introduced in the ECD do, perhaps, as expected simply on the basis of distance. As was the case for the wild-type human α7-AChR, the Hill coefficient for MLA turned out to be ∼1 for the two chimeric constructs, thus suggesting that the neurotransmitter-binding sites remained identical and independent despite the mutations.

The structure of the α7-AChR bound to MLA has not yet been solved, and MLA is a larger molecule than both epibatidine and EVP-6124 (Fig. 11). Therefore, we cannot rule out the possibility that the different MLA affinities of the human and chicken α7-AChR’s ECDs (Fig. 17 A) are due to close contacts between MLA and one or more of the 13 amino acids that differ between these two orthologs. Of these, residue 172 (residue 149 in the alternative numbering system used in (Noviello et al., 2021)) approaches epibatidine the closest (4.1 Å in PDB ID code 7KOX; (Noviello et al., 2021)), whereas residue 56 approaches EVP-6124 the closest (3.1 Å in PDB ID code: 7EKP; (Zhao et al., 2021)); all other 11 residues lie farther than 5 Å from these bound agonists. Hence, we mutated only these two residues of the human–*C. elegans* chimera (Ser-56 and Ser-172) to their chicken counterparts (both threonine) to estimate the degree to which the different MLA affinities of the human and chicken α7-AChR’s ECDs may be attributed to these closer-to-the-orthosteric-site substitutions. MLA–toxin competition curves for this double mutant revealed that the half-competition concentration only changed from 68 ± 6 nM in the human–*C. elegans* chimera to 32 ± 4 nM in the S56T + S172T mutant, whereas that of the chicken–*C. elegans* chimera is 4.9 ± 0.5 nM (Fig. 17 A and Table 1). Thus, it follows that it is the other 11 chicken-*versus*-human amino-acid substitutions that account for most of the difference between the MLA closed-state affinities of these two α7-AChR orthologs.

The notion that changes in the amino-acid sequence of the TMD have a comparatively minor effect on the ligand-binding properties of the α7-AChR ECD, inferred above from the observations with MLA, can very likely be generalized to all orthosteric ligands. Thus, the differences observed between the competition curves of the human α7-AChR and the human–*C. elegans* α7-AChR–β-GluCl chimera with the agonists nicotine and carbamylcholine (Fig. 17, B and C) may largely be ascribed to differences in these channels’ unliganded-gating equilibrium constants. We found the chimera’s curves to be right-shifted relative to those of the α7-AChR, regardless of the competing agonist, consistent with the chimera having a lower-than-wild-type unliganded-gating equilibrium constant (Fig. 4 A; Fig. 5 A; and Table 1). Also consistent with this notion is the lower Hill coefficients obtained for this chimera, both for nicotine and carbamylcholine (Fig. 6 A and Table 1).

The toxin–agonist competition curves obtained with the chicken–*C. elegans* chimera (Fig. 17, B and C), on the other hand—left-shifted for nicotine and nearly overlapping for carbamylcholine relative to those obtained with its human–*C. elegans* counterpart—exemplify the difficulty in comparing the behavior of pairs of receptors when both unliganded gating and ligand affinities are expected to be different. Indeed, although these chimeras share the same TMD, the 13 amino-acid differences at the ECD are also likely to have caused changes in the equilibrium constant of unliganded gating. The effect of these mutations on the unliganded-gating equilibrium constant is, of course, the same irrespective of the agonist used, but the effects on agonist affinities are likely to be ligand-specific.

## DISCUSSION

Although probing the function of ion channels without measuring the transport of ions through them may seem oxymoronic, a number of experimental situations (see Introduction) call for such indirect approaches. Here, working on members of the Cys-loop-receptor superfamily of ligand-gated ion channels (pLGICs), we focused on the practical implementation of, and the interpretation of results from, ligand-binding assays. To some extent, it could be said that the latter are to ligand-gated ion channels what gating-current recordings are to voltage-dependent channels: a means to probe the function of a domain that, at least in wild-type channels, is coupled to the channel’s activation gate. An important difference, however, is that ligand-binding studies do not require that the two ends of the ion channel face electrically separate compartments, and thus, binding assays can also be applied to detergent-solubilized or nanodisc-reconstituted receptors.

Ligand-binding studies of pLGICs are, by no means, new (Fulpius et al., 1972; Maelicke et al., 1977; Miledi et al., 1971; Miledi and Potter, 1971; Weber and Changeux, 1974a, 1974b, 1974c). This is the reason why we were surprised to note a lack of uniformity in published protocols, large disparities in the estimates obtained by different groups for the same parameter, and a general disregard for constraints placed on the experimental observations by simple theoretical considerations. We decided to pursue the equilibrium-type of ligand-binding assays rather than the (much-faster and more frequently-used) protection-type kinetic assays because the former seemed, overall, more straightforward. Indeed, we surmise that some of the inconsistencies associated with the application of the kinetic approach may have arisen from the (admittedly challenging) accurate estimation of only the initial rate of labeled-ligand binding.

As elaborated in the Results section, equilibrium binding-competition assays present difficulties, too. A major one is the fact that the interaction between the labeled ligand and the receptor (seldom of interest) gets in the way of the characterization of the interaction between the unlabeled ligand and the receptor. To address this point, we performed [^125^I]-α-BgTx saturation curves with every new α7-AChR construct so as to learn what (fixed) concentration of labeled toxin had to be used in competition assays. This is a crucial step that ensured that comparisons between different unlabeled ligands, different constructs, and different experimental conditions were unaffected by the properties of the toxin–channel complex. Using the same labeled ligand for all of our assays certainly helped mitigate this inconvenience. Another difficulty— particularly when using slowly dissociating ligands such as α-BgTx—is the need to make a judgement as to when the system is close enough to equilibrium. In our case, we incubated the reactions at different temperatures for different times and deemed them to have approached equilibrium to a satisfactory degree when the fitted empirical parameters changed little with longer incubations. We note that the strong effect of temperature on the kinetics of approach to equilibrium seems to have gone unnoticed in previous applications of this method. Indeed, raising the incubation temperature to 37°C sped up the reactions’ time courses considerably.

Undoubtedly, equilibrium assays of the sort we described here are too time-consuming and labor-intensive to be useful as tools for the high-throughput screening of drugs. Rather, they are meant to be used in the context of the detailed mechanistic characterization of receptor-channel operation, an integral aspect of the design of new drugs that should not be overlooked. In this latter regard, it is important to bear in mind that an abundance of functional—and more recently, structural—data point to the notion that pLGICs form a mechanistically homogeneous group of proteins. Hence, conclusions drawn from studies of α-BgTx-binding AChRs may well hold for the rest of the superfamily.

We would like to emphasize that the guidelines we provided here for the implementation and interpretation of concentration–response curves are valid for any antagonist or inverse agonist acting as the labeled ligand irrespective of their dissociation kinetics. Although assays that require the physical separation of bound from unbound label are most accurately performed with slowly dissociating labeled ligands, more recently developed technologies (for example, “scintillation-proximity assays”; SPA (Udenfriend et al., 1985) eliminate the need for this step, and thus open up the field to all other pLGICs for which slowly dissociating competitive ligands are not known or are difficult to obtain. Whether this faster type of assay (intended, essentially, to allow for the high-throughput screening of ligands) yield data of high-enough a quality to illuminate ion-channel mechanisms, however, remains to be ascertained.

For the sake of conciseness—and because, here, we used α-BgTx as the label—we did not elaborate on the mechanistic interpretation of concentration–response curves obtained from assays in which the labeled ligand is an agonist. However, several fast-dissociating pLGIC agonists are commercially available in radiolabeled form, and their use in equilibrium-type competition experiments has been increasing as the use of the SPA technology is becoming more widespread. A cursory theoretical analysis of the relationship between the empirical parameters of the corresponding ligand-binding curves and the underlying equilibrium constants of state interconversions reveals that, although some aspects remain the same regardless of whether the labeled ligand is an agonist, an antagonist or an inverse agonist, others differ in important ways. For example, among the latter, half-saturation concentrations of labeled agonists are not dissociation equilibrium constants (*K_D_* values) from the closed state, and Hill-coefficient values from competition curves are highly sensitive to the ratio between the fixed and half-saturation concentrations of the labeled ligand. Clearly, as the use of agonist labeled ligands in equilibrium-type competition experiments increases, so does the need of enhancing the rigor and attention to theoretical detail with which these quantitative methods are applied. This is especially true if the obtained numbers are more than just mere numbers, and instead, are expected to help us understand how ligand-gated ion channels work.

## ACKNOWLEDGMENTS

We thank S. Gough for experiments performed during the initial stages of this project, and Y. Paas (Bar-Ilan University, Ramat Gan, Israel), W. N. Green (University of Chicago, IL), and S. M. Sine (Mayo Clinic College of Medicine, MN) for cDNAs. This work was supported by a grant from the US National Institutes of Health (R01-NS042169 to C.G.).

The authors declare no competing financial interests.

## Author contributions

N. Godellas: Conceptualization, formal analysis, investigation, methodology, visualization, writing – original draft, writing – review & editing. C. Grosman: Conceptualization, funding acquisition, project administration, supervision, writing – original draft, writing – review & editing.

